# Early-life *Helicobacter pylori* infection in mice alters endocrine signaling and accelerates adipose expansion during high-fat feeding

**DOI:** 10.1101/2025.04.27.650497

**Authors:** Sigri Kløve, Katrine B. Graversen, Jacob A. Rasmussen, Juliana Assis, Jonas Schluter, Martin J. Blaser, Sandra B. Andersen

## Abstract

Perturbations to early-life microbial colonization can shape immune and metabolic development, predisposing a host to obesity. We hypothesized that neonatal infection with the ancient human symbiont *Helicobacter pylori* exacerbates diet-induced metabolic responses and is accompanied by alterations in endocrine and inflammatory regulation. To test this, C57BL/6JRj neonatal mice were infected with *H. pylori* or left uninfected and, after weaning, exposed to a high-fat diet in a long-term (5-month) study of microbiome composition and a short-term (3-week) study of circulating biomarkers and microbiome composition. In the short-term intervention, infected mice showed increased visceral adiposity. Infection was associated with altered ghrelin and leptin responses across fasting conditions, accompanied by elevated MCP-1 and IL-6, particularly in males. *H. pylori* infection also reinforced diet-induced shifts in gastrointestinal microbiome composition. In the long-term study, there were no differences in adiposity between infected and control mice, suggesting that diet was the dominant determinant of adiposity at this stage. Together, our findings suggest that early-life *H. pylori* infection amplifies the initial endocrine, inflammatory, and adipose responses to a high-fat diet.

## INTRODUCTION

Obesity and related disorders are increasing globally, alongside shifts in gut microbiome composition that may contribute to disease risk ^1^. Microbial colonization begins at birth, and early-life perturbations of the microbiota are associated with increased risk of overweight and obesity ^2–4^. These disruptions can alter immune system development during a critical early window ^5^, potentially promoting less tolerant, more reactive immune responses and increasing susceptibility to allergic, autoimmune, and inflammatory diseases ^6^. Given the lasting impact of early-life microbiota perturbations, attention has turned to “ancestral microbes” with long-standing host associations. *Helicobacter pylori* is one such bacterium, colonizing humans for at least 300,000 years ^7–9^ and typically transmitted from mother to child ^10^.

Our co-evolution with *H. pylori* has produced a complex relationship: while most carriers remain asymptomatic, ∼10% develop gastroduodenal diseases, including ulcers and cancer ^11^. Conversely, *H. pylori* has been linked to protection against conditions such as gastroesophageal reflux disease, childhood asthma, and allergies ^12,13^, diseases all increasing in incidence as *H. pylori* has been disappearing. Outcomes may be affected by factors such as age at acquisition ^14,15^, family structure ^16^, and host–microbe genomic compatibility ^17^. Although still present in ∼50% of the global population, prevalence is declining particularly in industrialized countries due to improved sanitation, antibiotics, and eradication efforts ^18,19^. Understanding the consequences of failing to acquire *H. pylori* early in life may provide insight into how the absence of ancestral microbes shapes disease risk in modern environments.

A major public health challenge in industrialized societies is the rising prevalence of obesity^20^. An inverse correlation between *H. pylori* prevalence and obesity across countries ^21^ has prompted investigation into its potential role in metabolic regulation. *H. pylori* may influence host metabolism through multiple pathways, including its modulation of hormone production, inflammation, and the gut microbiome ^22,23^. The hormones ghrelin and leptin are central to energy balance: ghrelin, largely produced in the stomach, regulates hunger and has both pro- and anti-inflammatory effects ^24^, while leptin, primarily secreted by adipose tissue, links endocrine and immune function, by promoting inflammatory cytokine production ^25,26^. *H. pylori* colonization has been associated with reduced ghrelin levels and variable leptin responses in humans ^27–33^. In mice, early-life *H. pylori* infection is characterised by a systemic anti-inflammatory response in addition to local inflammation ^12,14,34^. This may affect the development of obesity, which is characterised by low-grade inflammation in adipose tissues ^35^.

We recently reported a strong positive association between *H. pylori* seropositivity and hyperglycemia, but not obesity, in Danish children and adolescents ^36^, suggesting a metabolism-modulating role of early-life infection. However, both increased and decreased obesity risk have been linked to *H. pylori* ^37,38^, potentially reflecting confounding by factors such as ethnicity, socioeconomic status, age, and sex ^39,40^. To better isolate causal effects, we used a mouse model to examine whether early-life *H. pylori* infection influences metabolism under diet-induced obesity. While the impact of a Western high-fat diet (HFD) has been studied in adult-infected mice ^41,42^, early-life infection remains less explored. Here, we show that early-life *H. pylori* infection, combined with HFD, promotes visceral fat accumulation and alters appetite-regulating hormones, inflammatory cytokines, and gut microbiome composition in a sex-specific manner.

## METHODS

### Animal experiments

Breeding pairs (two females, one male) of C57BL/6JRj mice (Janvier, France) were established. Neonates received ∼50 μL of 10^9^ CFU *H. pylori* PMSS1 or sterile media by oral gavage at 6 and 7 days of age (**Fig. 1**). When both females produced litters, pups were split across treatments. Mice were weaned at day 25 and randomized (3–4 per cage) by sex and treatment, with litters mixed within groups. Two complementary experiments were performed: (1) a long-term study designed to examine the consequences of neonatal *H. pylori* infection following prolonged exposure to an obesogenic diet, with primary outcomes being gastrointestinal microbiome composition and adipose tissue phenotype. All mice received a HFD (Research Diets D12331; 58 kcal% fat, 25 kcal% carbohydrate, including sucrose) one week post-weaning and maintained for 20–21 weeks (**Fig. 1A**). (2) a short-term experiment designed to investigate how infection affects the early endocrine, inflammatory, and microbiome responses to HFD, including both HFD and control diet groups (CD; Research Diets D12328; 10 kcal% fat, 73 kcal% carbohydrate, primarily cornstarch) over a three-week intervention starting two weeks post-weaning (**Fig. 1C**). The purified diets differ primarily in fat content but also in carbohydrate composition. Diet initiation differed between experiments (one versus two weeks post-weaning) due to a delay in diet delivery. Target group sizes were >5 animals (30% detectable difference; α = 0.05, power = 0.80), but groups were unbalanced (3–9 animals) due to pre-sexing treatment allocation. Bayesian regression analysis was used to estimate treatment effects and characterize the associated uncertainty in the presence of small, unequal group sizes. Primary conclusions are based on groups of 5–9 animals from 3– 4 litters across 2–3 cages, limiting litter and cage effects (**Table S1**). Experiments were conducted ∼1 year apart in separate rooms within the same facility. Mice were housed in ventilated cages at 22 ± 1 °C under a 12 h light–dark cycle with ad libitum food and water. All procedures were approved by the Danish Animal Experimentation Inspectorate (2020-15- 0201-00432).

**Figure 1:**
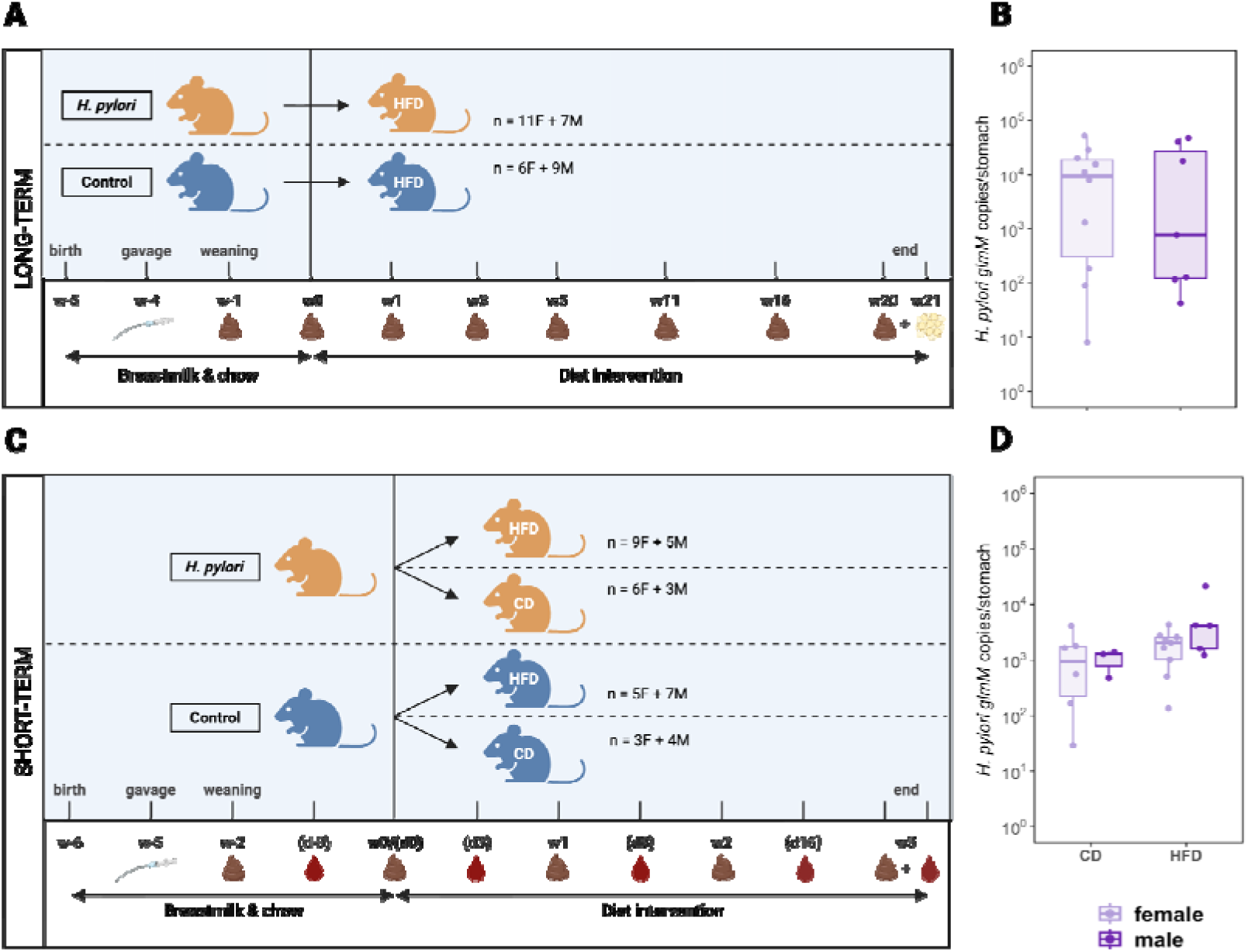
Experimental overview and *Helicobacter pylori* colonization levels. **A** Schematic overview of the long-term experiment. C57BL/6JRj mice were orally gavaged with *H. pylori* PMSS1 (n = 18) or sterile media (control; n = 15) at postnatal days 6–7. One week after weaning, all mice were changed to a high-fat diet (HFD; 58% kcal fat with sucrose). Body weight and fecal samples were collected longitudinally. Mice were sacrificed 20 (females) or 21 (males) weeks after diet initiation for tissue collection (perigonadal white adipose tissue, stomach tissue, ileum, cecum, and colon content). **B** Gastric *H. pylori* colonization levels (copy number / ng total DNA) at termination of the long-term experiment. No clear differences between sexes were observed. **C** Schematic overview of the short-term experiment. Mice were gavaged as above (n = 23 infected, n = 19 control). Two weeks after weaning, mice were assigned to HFD or control diet (CD; 11% kcal fat). Body weight, fecal samples, and plasma were collected longitudinally. Mice were sacrificed three weeks after diet initiation. **D** Gastric *H. pylori* colonization levels at termination of the short-term experiment. No clear differences between diets or sexes were observed. Sampling times are noted with a w for week and d for day. Time is relative to diet intervention, which was initiated one week apart in the two experiments, hence the differences in birth, gavage and weaning time nomenclature between A and C.

### *H. pylori* PMSS1 infection

The *H. pylori* strain PMSS1 ^14^ was grown on TSA Sheep Blood plates (Thermo Fisher Scientific) for 72h at 37 °C, 10% CO_2_ and 6% O_2_. Bacteria were transferred to liquid Brucella broth (BD Biosciences) with 10% heat-inactivated fetal bovine serum (FBS, Sigma-Aldrich) and 0.06 mg/ml Vancomycin (Sigma-Aldrich) and incubated overnight with shaking at 100 rpm. Cultures were diluted with fresh media to an OD_600_ of 0.1, shaken for 2-3 h, and centrifuged at 900 g at room temperature for 10 min. The bacterial pellet was resuspended in Brucella broth with 10% h FBS. Serial dilutions were inoculated for CFU counting.

### Experimental procedures

Body weight was recorded weekly from diet initiation. Fecal pellets were collected at selected timepoints (**Fig. 1**) by placing animals in a beaker until defecation; pellets were collected with tweezers, snap-frozen on dry ice, and stored at −70 °C. Equipment was cleaned between animals. In the short-term experiment, mice were fasted for 4 h in the morning. Under isoflurane anesthesia, blood was collected from the submandibular vein into pre-cooled K2EDTA tubes (BD Biosciences), immediately supplemented with protease inhibitor (1×) and DPP-IV inhibitor (0.1 mM), kept on ice, centrifuged (2000 g, 4 °C, 10 min), and plasma stored at −70 °C.

### Necropsy procedures

Mice were fasted overnight (12 h), anesthetized with isoflurane, and euthanized by cervical dislocation. Blood was collected by cardiac puncture and processed as described above. The forestomach was removed, and the glandular stomach was divided longitudinally into four equal sections for standardized downstream analyses. Luminal contents from the ileum, cecum (long-term experiment only), and colon were collected, snap-frozen on dry ice, and stored at −70 °C.

### Isolation, cell counting and flow cytometry of stromal vascular cells (SVC) from perigonadal white adipose tissue (WAT)

In the short-term experiment, SVCs were isolated from the right perigonadal WAT (males) or both sides (females) and processed as described ^43^. Cells were counted [Countess 3 (Invitrogen)] and 0.25 - 1 × 10^6^ cells/well were seeded in a 96 U-bottom well plate (Corning Costar), and stained for surface markers as described ^43^ (see also SOM methods).

### Measurement of metabolic biomarkers

In the short-term experiment, plasma levels of 10 metabolic markers were quantified using a customized U-PLEX Meso Scale Discovery (MSD) assay following the manufacturer’s protocol. The panel included the obesity panel validated by the manufacturer, covering: total ghrelin, leptin, insulin, C-peptide, total peptide YY (PYY), total glucagon-like peptide-1 (GLP-1), and added interleukin 10 (IL-10), interleukin 6 (IL-6), tumor necrosis factor α (TNF-α) and monocyte chemoattractant protein 1 (MCP-1), selected based on relevance to *H. pylori* infection ^44–48^ and obesity ^49^. Plates were read on a MESO QuickPlex SQ 120 instrument (MSD). Fasting glucose levels were measured but are not reported because isoflurane anesthesia during sampling was recognized to confound the measurements.

### *H. pylori* PMSS1 quantitation

*H. pylori* colonization was quantified from DNA extracted from one quarter of the glandular stomach using the DNeasy Blood & Tissue Kit (Qiagen), with overnight lysis (56 °C, 300 rpm) and elution in 100 μL. Infection was first confirmed by conventional PCR. Positive samples were quantified by absolute qPCR using a standard curve in 20 μL reactions run in duplicate. The standard curve was generated from serial dilutions of a plasmid containing the target region, and copy numbers were calculated accordingly. *H. pylori* levels were normalized to total DNA per sample, measured by Qubit dsDNA high-sensitivity assay (Invitrogen), as a proxy for tissue input (see also SOM methods).

### DNA extraction, 16S metabarcoding library preparation and bioinformatical analyses

DNA was extracted from fecal pellets and intestinal contents (ileum, cecum, colon) using the ZymoBIOMICS Quick-DNA Fecal/Soil Microbe 96 Magbead Kit (Zymo Research) in randomized batches, including extraction controls. The V3–V4 region of the 16S rRNA gene was amplified with 341F/805R primers, with positive and negative controls included on each plate. Amplicons were purified, indexed using Nextera XT (Illumina), pooled, and sequenced on an Illumina MiSeq platform. Raw sequencing data were demultiplexed and converted to FASTQ using Illumina bcl2fastq2 (v2.20). Subsequent processing was performed with DADA2 ^50^, including quality filtering, trimming, read merging (minimum 12 bp overlap), chimera removal, and ASV inference. Taxonomy was assigned to genus level using the SILVA v138 database. Post-clustering (98% similarity) was performed with *DECIPHER* ^51^ to retain resolution of mock community strains. Contaminants and well-to-well leakage were identified and removed using SCRuB ^52^ based on extraction and non-template controls. Eukaryotic ASVs were excluded prior to analysis (see also SOM methods).

### Statistical analyses

All analyses were performed in R (v4.3.3) and RStudio (v.2025.09.0+387). Bayesian regression models were fitted using *brms* ^53^ with post-processing in *tidybayes* ^54^. Weakly informative priors were used unless stated otherwise. Models were estimated via Hamiltonian Monte Carlo (4 chains, 4,000–6,000 iterations, including 2,000–3,000 warm-up). Appropriate likelihoods were specified (Student-t for continuous outcomes, negative binomial for counts, beta for bounded responses). Inference was based on posterior medians, 80% and 95% credible intervals, and posterior probabilities of effects > 0. Results are reported as estimated effects with corresponding 95% credible intervals and posterior probabilities. Effects were interpreted as strong when 95% intervals excluded zero, and as tendencies when 80% intervals excluded zero. Models included ANOVA-style main effects and interactions, with simple effects used for interpretation. Effect sizes are reported as posterior median differences. Additional packages included *vegan* ^55^, *tidyverse* ^56^, *speedyseq* ^57^, *reticulate* ^58^, *emmeans* ^59^, and *fantaxtic* ^60^. Visualizations were generated using *ggplot2* ^61^, *cowplot* ^62^, *patchwork* ^63^ and *ggpubr* ^64^. Full model code is provided in the data availability section.

#### H. pylori quantitation

*H. pylori* colonization levels were compared between experimental groups and conditions with log_10_-transformed standardized bacterial load as the outcome and sex and diet as predictors.

#### Body and perigonadal WAT weight

Body weight trajectories were analyzed with time (weeks) as a continuous predictor, allowing group-specific differences in trajectories. Random intercepts and random slopes for time were included for each mouse to account for repeated measurements. Time was centered at the mean study week. WAT weight measured at termination was analyzed using a linear model. One control male receiving a HFD in the short-term experiment had missing WAT data.

#### Cytokine and hormone assay

Prior to diet intervention, log_10_-transformed biomarker levels were analyzed with treatment and sex as predictors. Post-diet models additionally included diet and time (categorical) to capture non-linear trends, and mouse ID was included as a random intercept to account for repeated measures. C-peptide was excluded due to values below detection limits; for other biomarkers, values below detection were set to the lower limit of the standard curve before log-transformation. In total, 63 measurements were below the assay detection limit or otherwise unavailable, the majority of which (51) were IL-6 measurements from *H. pylori* female mice (HFD: 13; CD: 16). Although individual values may reflect either technical failure or very low cytokine concentrations, their strong enrichment in these females is consistent with generally lower circulating IL-6 levels in this group. Associations between biomarkers, *H. pylori* colonization, and adiposity were analyzed in infected animals only. Models included sex, diet, standardized *H. pylori* load, and standardized WAT weight. Model comparison (LOO-CVl ^65^) provided limited support for interactions with WAT weight, which was therefore retained as a main effect, while *H. pylori* effects were allowed to vary by sex and diet. Posterior predictions were evaluated across observed ranges of *H. pylori* load and WAT weight, holding the alternate predictor at its mean. For ease of interpretation, effect estimates reported in the text are presented as fold increases or percentage decreases.

#### Flow Cytometry

Flow cytometry data were gated and analyzed separately for each sex. The log_10_-transformed F4/80^+^ CD11b^+^ counts per gram perigonadal WAT, the proportion of CD11c^+^ M1-like macrophages, CD301^+^ M2-like macrophages, and the ratio of CD11c^+^ to CD301^+^ macrophages was analysed with treatment, diet, sex, and their interactions.

#### Alpha Diversity

Alpha diversity metrics from longitudinal fecal samples were analyzed, with Shannon diversity modeled with a Gaussian likelihood, and Observed Species using a negative binomial distribution. Mouse ID was included as a random intercept to account for repeated measures prior to diet switch. Gastrointestinal samples collected at a single time point were analyzed using equivalent models without random effects.

#### Beta Diversity

Longitudinal changes in fecal microbiota were assessed using within-mouse Bray–Curtis dissimilarity. Calculated distances were restricted to within-individual pairs, and modeled as a function of standardized time between samples. Pairwise dissimilarities (0–1) were analyzed using beta regression (logit link) with time separation and a diet × treatment × sex interaction, including random intercepts and slopes for each mouse. Posterior expected dissimilarities were estimated at zero time separation to enable comparisons across conditions, with infection effects summarized within each sex–diet combination. For terminal gastrointestinal samples, PCoA axis scores were analyzed using multivariate Gaussian regression with correlated residuals. Effect estimates represent posterior differences between group centroids, capturing multivariate community separation.

#### Microbiome composition

Microbiome data were processed using *phyloseq* ^66^ and subset into pre-diet, post-diet, and gastrointestinal sample groups. Taxa were aggregated at the family level. In the LT experiment, post-diet family-level data were filtered before transformation to retain families present in at least 10% of samples and reaching a maximum relative abundance of at least 0.1% in at least one sample. To address zero values, a pseudocount equal to half the smallest non-zero relative abundance was added prior to renormalization. Data were transformed using the centered log-ratio (CLR) transformation using *compositions* ^67^ to account for compositional constraints, and the effects of diet, treatment, sex, and their interactions were estimated.

#### Muti-omics data integration

For *H. pylori* infected animals in the short-term experiment, an exploratory multi-omics factor analysis was performed using *MOFA2* ^68^ to identify shared sources of variation across datasets, including CLR-transformed colon microbiome composition of the nine most differentially abundant families, log_10_-transformed biomarker levels, WAT weight, and *H. pylori* load. All features were standardized and matched across samples. Models were fit using Gaussian likelihoods with variational inference. Latent factors were interpreted based on their associations with metadata and feature loadings.

## RESULTS

### Stable *H. pylori* PMSS1 gastric colonization of mice subjected to HFD or control diet

In the long-term experiment (HFD for ∼5 months from one-week post-weaning; **Fig. 1A**), gastric *H. pylori* levels ranged from 10^0^ to 10^5^ copies/ng DNA, with medians in the 10^3^-10^4^ range (**Fig. 1B**), with no sex difference (posterior median difference: -0.19, 95% CI [-1.08, 0.71], posterior probability 64%). In the short-term experiment (3 weeks on control or HFD; **Fig. 1C**), levels ranged from 10^1^ to 10^4^ copies/ng DNA with medians in the 10^3^ range (**Fig. 1D**) and varied substantially between individuals, with tendencies for higher loads in the animals receiving the HFD (females: 0.34, 95% CI [-0.21, 0.89], 88%; males: 0.56, 95% CI [-0.18, 1.29], 94%), and for higher loads in males than females receiving the HFD (0.39, 95% CI [-0.20, 1.00], 90%). There were no sex differences for mice receiving the control diet (0.16, 95% CI [-0.51, 0.88], 68%). Among HFD-fed mice, *H. pylori* loads were similar between experiments (females: -0.20, 95% CI [-0.97, 0.62], 68%; males: 0.23, 95% CI [-0.82, 1.27], 33%).

## SHORT-TERM EXPERIMENT

### Early-life *H. pylori* infection accelerates weight gain and perigonadal WAT accumulation

We found that weight gain varied by sex and context. Within the *H. pylori* infected males, HFD increased weight gain compared to control diet (Δ 1.88 g, 95% CI [0.15, 3.61], posterior probability = 98%), with a similar tendency in females (Δ 1.21 g, 95% CI [−0.07, 2.47], 97%; **Fig. S2**). However, there was no strong evidence for overall main effects of infection or diet, or their interaction on weight gain. In contrast, WAT weight showed a clear infection × diet interaction: *H. pylori* males on HFD had 85% higher WAT weight than controls, and females 57% higher (males: 95% CI [40.3, 146], females: 95% CI [7.27, 163], posterior probabilities >98%; **Fig. 2**). No effect of infection was observed in mice receiving the control diet, while HFD showed a borderline negative effect in control females.

**Figure 2:**
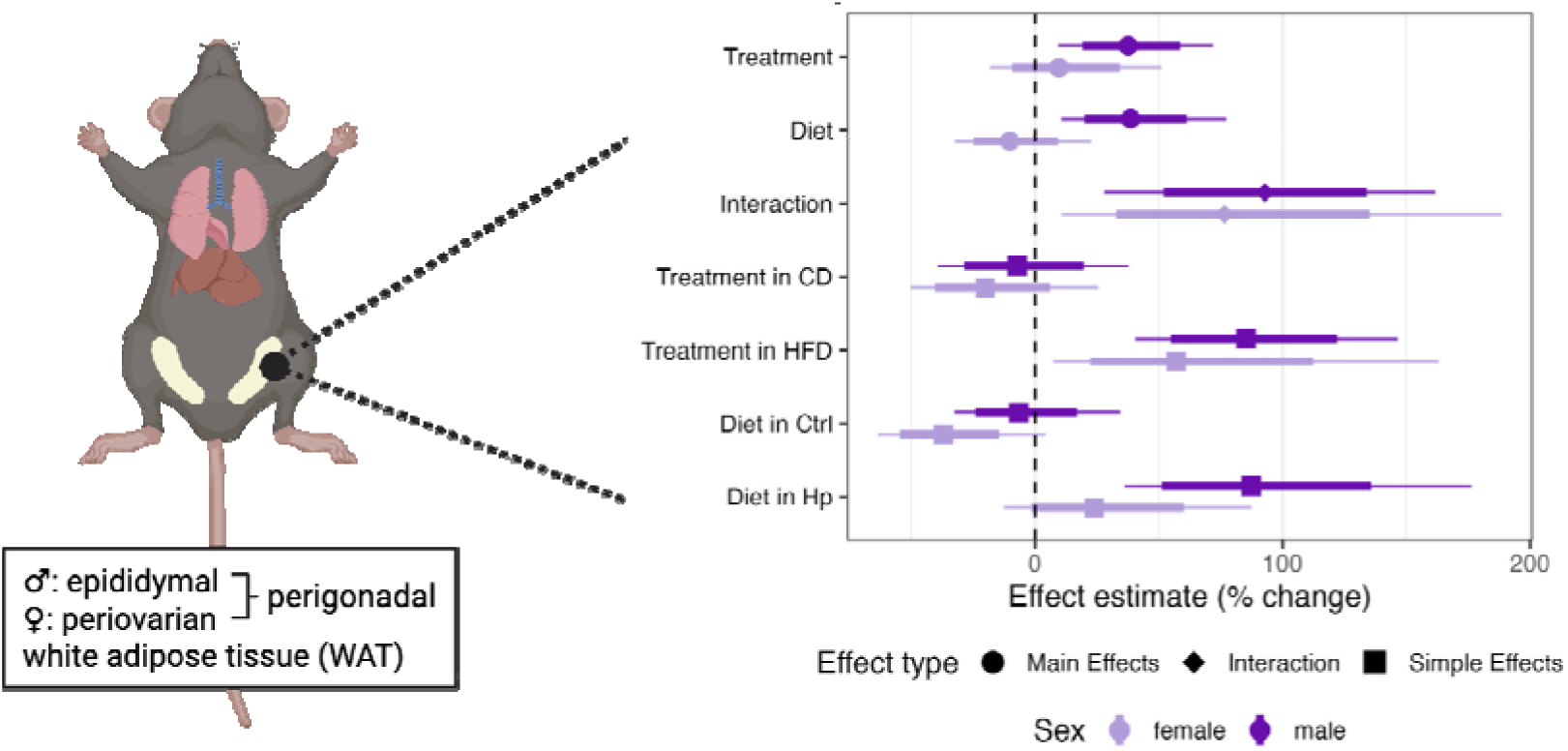
Analyses of perigonadal white adipose tissue (WAT) weight at termination of the short-term experiment. Bayesian linear regression revealed a diet × infection interaction, where *H. pylori*–infected males and females receiving a high-fat diet (HFD) had higher WAT weight compared with control animals on HFD. Forest plots showing posterior median effect estimates for main effects (treatment, diet, and their interaction; *H. pylori* vs control) and corresponding simple effects (treatment within each diet and diet within each treatment). Error bars show 95% credible intervals (thin lines) and 80% credible intervals (thick lines) for percentage change in WAT weight relative to the control group. Effect types are indicated by symbol shape (dot, main effects; diamond, interaction effects; square, simple effects), while light and dark purple denote females and males, respectively. The horizontal dashed line indicates no effect.

### *H. pylori* infection impacts circulating levels of metabolic hormones and cytokines

We collected blood samples before and after the diet intervention to assess selected metabolic biomarkers. At baseline, leptin levels were 2-fold higher in *H. pylori* males compared to controls (95% CI [1.2, 3.3], posterior probability: 100%). MCP-1 levels were ∼18% lower (95% CI [0.3, 32], 98%), while TNF-α showed a similar but weaker trend (∼24% lower, 95% CI [43% lower to 2% higher], 97%). Similar patterns were observed in females (**Fig. 3**).

**Figure 3:**
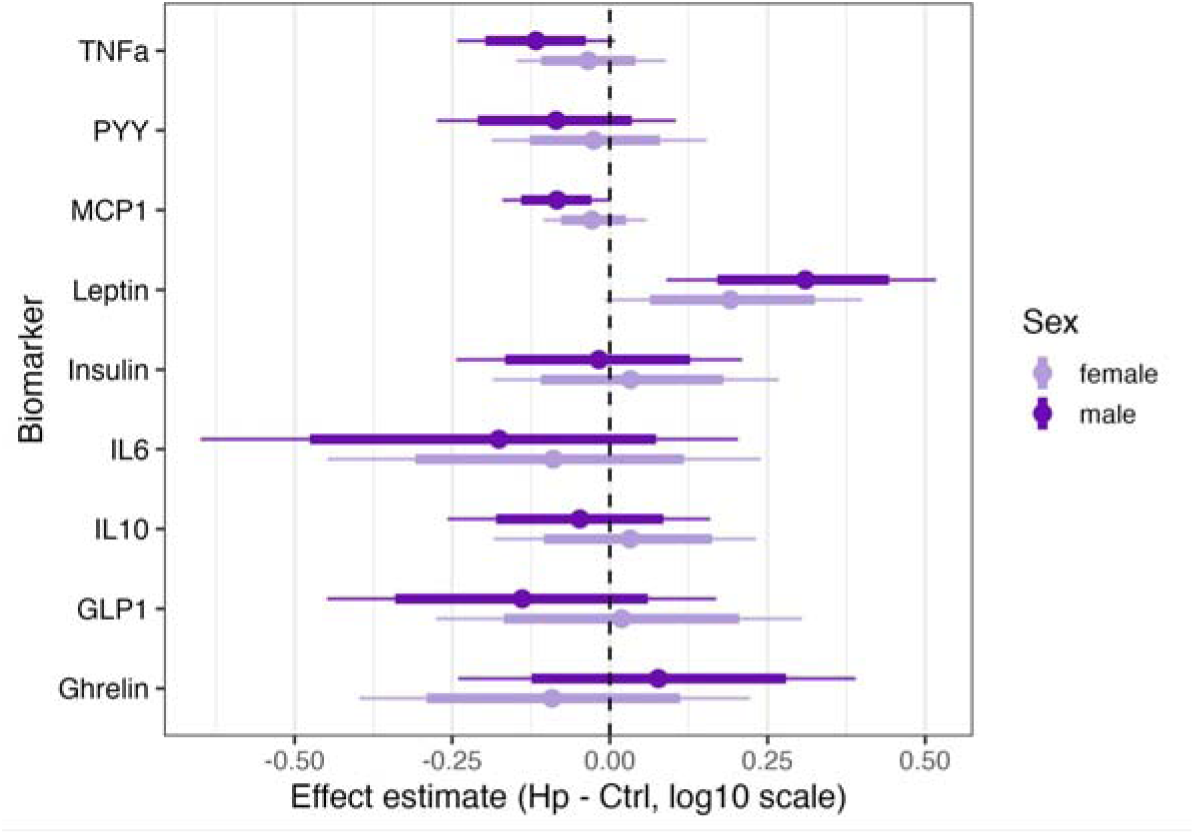
Differences in levels of nine biomarkers measured from serum samples with a Meso Scale Discovery kit, between *H. pylori*–infected and control mice sampled at baseline on a chow diet in the short-term experiment. Forest plots showing posterior median effect estimates for the effect of treatment (*H. pylori* vs control). Error bars show 95% credible intervals (thin lines) and 80% credible intervals (thick lines). An effect estimate of 0.25 corresponds to approximately a 1.78-fold change in biomarker concentration in *H. pylori*–infected animals compared with controls. Light and dark purple denote females and males, respectively. The horizontal dashed line indicates no effect.

During the diet intervention, biomarker profiles varied by *H. pylori* infection, diet, sex, and time. In males, *H. pylori* infection was associated with higher baseline ghrelin (∼2.1-fold; 95% CI [1.1, 4.2], 98%), which for the animals receiving the HFD increased when sampled after a 4-hour fast. Levels, however, failed to rise after a 12-hour fast and tended lower than that of control males at termination (**Fig. 4, S3A**). Leptin remained ∼2-fold higher in infected males and did not decline with extended fasting, resulting in markedly higher levels than controls (8.7-fold; 95% CI [3.7, 20.0], 100%; **Fig. 4, S3A**). IL-6 was initially lower (∼76%), but increased strongly after 12-hour fasting in HFD-fed infected males (8.5-fold; 95% CI [3.3, 20.7], 100%), while decreasing in controls (**Fig. 4, S3A**). MCP-1 showed a similar pattern, shifting from borderline lower during HFD to higher after 12-hour fasting (1.8-fold; 95% CI [1.2, 2.7], 99%; **Fig. 4, S3A**). Insulin was elevated early (1.7-fold; 95% CI [1.0, 3.0], 98%), with a tendency for higher GLP-1 (**Fig. 4**).

**Figure 4:**
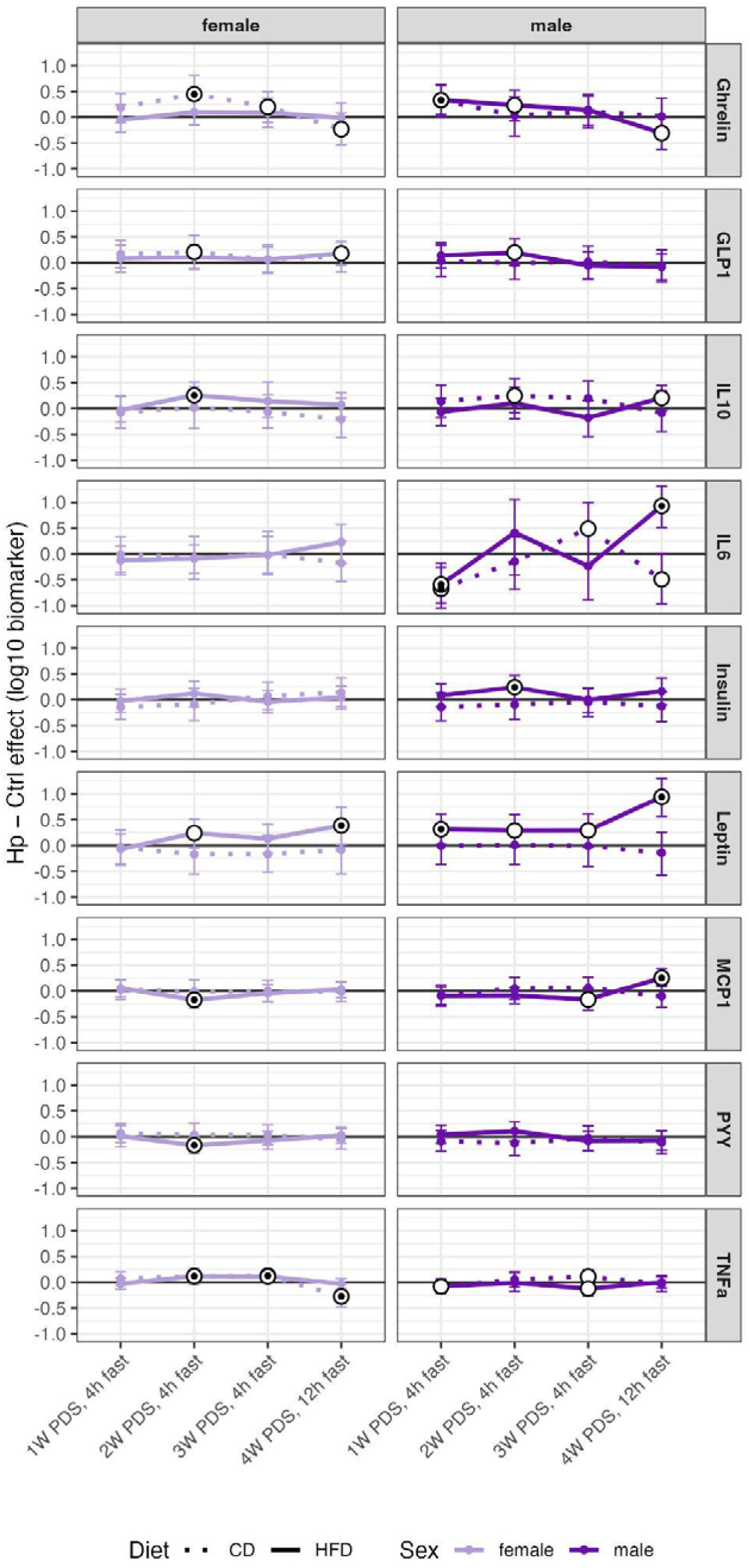
Trajectories of differences in levels of nine biomarkers comparing *H. pylori*–infected and control animals sampled one to four weeks after switching to a high-fat diet (HFD) or control diet (CD). The y-axis shows the effect size as the posterior median difference in biomarker concentration between treatments, expressed in log_10_ units. These differences represent the simple effects of treatment within a diet estimated from the full Bayesian mixed-effects model including treatment, diet, sex, and timepoint interactions, with random intercept for mouse ID to account for repeated measurements. An effect estimate of 0.5 corresponds to approximately a 3.2- fold change in biomarker concentration in *H. pylori*–infected animals compared with controls. As the plots depict the differences between the groups, a greater difference may reflect either an increase in levels of the *H. pylori* infected animals, or a decrease in levels of control animals. For the actual trajectories see Fig. S3. Error bars represent 95% credible intervals. White circles indicate estimates where the 80% credible interval excluded zero, and black dots within indicate estimates where the 95% credible interval excluded zero. For samples collected one to three weeks post diet switch (PDS), animals were fasted for 4 h; for the final time point animals were fasted for 12 h. Dotted lines denote animals receiving the CD and solid lines denote animals receiving the HFD. Light and dark purple indicate females and males, respectively.

In females, effects were most evident two weeks post-diet switch. Under HFD, *H. pylori* was associated with higher levels of IL-10 (1.8-fold) and TNF-α (1.3-fold), and lower MCP-1 and PYY (∼32%; >98%; **Fig. 4**). Ghrelin was higher under control diet (2.8-fold; 95% CI [1.2, 6.6], 99%; **Fig. 4**), with a tendency for increased GLP-1. TNF-α remained elevated under HFD, whereas control-fed infected females showed lower TNF-α at termination (−47%; 95% CI [6, 67], 100%; **Fig. 4**). GLP-1 tended to be higher under HFD at termination. Overall, infected females on HFD exhibited elevated leptin, IL-10, MCP-1, and TNF-α (**Fig. 4, S4**).

To investigate potential drivers of the altered hormone and cytokine levels at termination of the experiment after the 12-hour fast, we evaluated their associations with perigonadal WAT weight and *H. pylori* load in infected animals. WAT weight was positively associated with IL-6, leptin, and TNF-α, with weaker associations for IL-10 (positive) and GLP-1/PYY (negative; **Fig. S5**). In contrast, *H. pylori* load showed a more specific association with MCP- 1, particularly in HFD-fed males.

### *H. pylori* infection enhances HFD-induced macrophage changes in males

Obesity is marked by visceral WAT expansion and adipose tissue macrophage infiltration, contributing to the associated low-grade inflammatory state ^69^. We quantified macrophages in perigonadal WAT and observed a tendency toward 1.5-fold higher F4/80 CD11b counts per WAT weight in *H. pylori*–infected males on HFD compared to controls (posterior probability 92%; **Fig. S6A**). CD11c^+^ (M1 pro-inflammatory-like) macrophages increased with HFD, with the largest shift in *H. pylori* males (+2.36 vs. +1.75 percentage points in controls; ∼100%; **Fig. S6B**). A similar but weaker tendency was observed in females (+1.14 vs. +0.75; 92%). Overall, HFD effects on macrophage polarization were most pronounced in *H. pylori* males, consistent with fat accumulation and biomarker patterns. No clear effects were observed for M2-like macrophages (**Fig. S6C**).

### *H. pylori* infection primarily affects the gastric microbiome

As both HFD and *H. pylori* are known to alter the gastrointestinal microbiome composition ^22,70,71^, we examined potential combined effects using longitudinal fecal and terminal gastrointestinal samples. Prior to the dietary intervention, alpha diversity (observed species, Shannon index) was similar between groups (posterior probabilities <83%), and treatment did not influence the diet-induced decline in diversity (<89%). At weaning, *H. pylori* mice showed higher CLR abundance of Marinifilaceae in both sexes, which further increased after one week (male: 1.327, 95% CI [0.0691, 2.538], female: 0.933, 95% CI [0.0415, 1.885], >97.5%). Lachnospiraceae was higher in infected males (0.75, 95% CI [0.00690, 1.45], >97.5%) with the same tendency for females (0.410, 95% CI [-0.149, 0.971], 93%), while Lactobacillaceae tended to be lower in infected males (-0.585, 95% CI [-1.324, 0.156], 94%). After one week on HFD diet, infected females showed tendencies toward lower Clostridiaceae (-0.645, 95% CI [-1.563, 0.228], 92%), and higher Rikenellaceae CLR abundance (0.288, 95% CI [-0.140, 0.711], 90%), whereas infected males tended to have lower Prevotellaceae (-0.738, 95% CI [1.669, 0.172], 94%). By three weeks post-diet switch, microbiome composition was dominated by diet-associated shifts, with no substantial treatment effects detected.

At termination, the strongest treatment effects were observed in the gastric microbiome. Shannon diversity was lower in *H. pylori*–infected mice receiving the HFD (male: −2.07, 95% CI [−2.80, −1.30], female: −0.93, 95% CI [−1.62, −0.28], >97.5%; **Fig. S7**), and inversely associated with *H. pylori* load (slope −1.30, 95% CI [−1.80, −0.77], 99%). PCoA of Bray–Curtis distances showed increased community separation with infection (**Fig. S8**). Not surprisingly, *H. pylori* load was strongly positively associated with Helicobacteraceae abundance (slope 0.42, 95% CI [0.24, 0.59], 99%). Within infected animals, HFD increased Helicobacteraceae CLR abundance (male: 2.05, 95% CI [0.68, 3.57], female: 1.61, 95% CI [0.15, 3.09], >98%; **Fig. S9**), despite similar levels across groups. Infected males also showed lower Akkermansiaceae in stomach and colon (**Figs. S9–S10**). In the colon, infection amplified HFD effects, reducing Prevotellaceae and increasing Streptococcaceae, Lachnospiraceae, and *Eub. coprostanoligenes* group (**Fig. S10**). Helicobacteraceae was detected only in infected stomachs, except for four low-abundance samples (0.03–0.2%; **Table S2**), likely due to cross-contamination.

In an exploratory multi-omics factor analysis of infected animals, we identified a single dominant latent factor linking colonic microbiome composition, fasting biomarkers, adiposity, and *H. pylori* load. This factor was positively associated with *H. pylori* load in HFD-fed animals (median β = 0.78, 95% CI [0.21, 1.29], 99%), whereas evidence for an association in control diet animals was weak (**Fig. 5A**). Factor 1 also showed a positive association with adiposity across diet groups (median β = 0.37, 95% CI [0, 0.72], 95%), with little evidence for a diet-specific interaction (**Fig. 5B**). Factor 1 values differed strongly by diet, with higher scores in HFD-fed animals (median β = 1.56, 95% CI [1.11, 1.99], ∼100%; **Fig. 5C**). The factor was characterized by higher inflammatory (IL-6, TNF-α, MCP-1) and metabolic (leptin, insulin) markers, lower ghrelin, and microbiome shifts including increased Lachnospiraceae and Streptococcaceae and decreased Marinifilaceae and Prevotellaceae (**Fig. 5D**). These findings suggest that adiposity broadly shapes biomarker profiles along a shared diet-associated inflammatory–metabolic axis, with variation in *H. pylori* load associated with this host–microbiome axis among HFD-fed animals.

**Figure 5:**
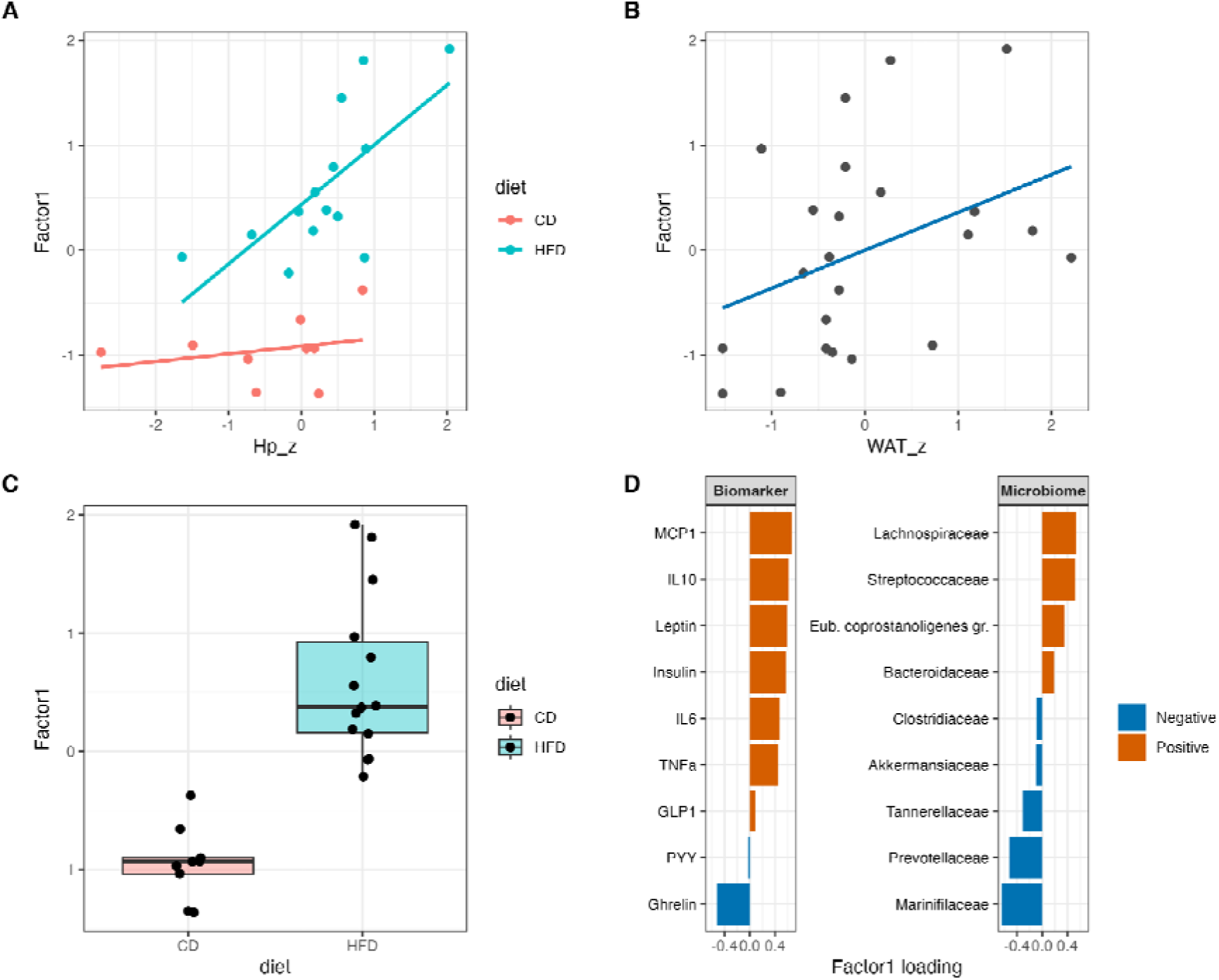
Multi-omics factor analysis identifies a coordinated inflammatory–metabolic axis in *H. pylori*–infected animals. Factor 1 represents the dominant latent pattern of coordinated variation shared across microbiome composition, circulating biomarkers, and host metabolic traits, consistent with an inflammatory–metabolic axis associated with adiposity and *H. pylori* burden. Analyses were performed in *H. pylori*–infected animals only. **A** Association between Factor 1 and *H. pylori* load (z-score). Each point represents an individual animal, coloured by diet (control diet, CD; high-fat diet, HFD). Lines indicate linear model fits. **B** Association between Factor 1 and adiposity, measured as standardized perigonadal white adipose tissue (WAT) weight (z-score). Each point represents an individual animal; the line indicates a linear model fit. **C** Distribution of Factor 1 values by diet (CD vs HFD). Points represent individual animals overlaid on boxplots (median and interquartile range). **D** Top feature loadings contributing to Factor 1 across microbiome family composition (CLR-transformed) and circulating biomarkers (log_10_-transformed). Loading magnitude reflects the strength of each feature’s contribution to Factor 1, with larger absolute values indicating stronger associations with the latent factor. Bars represent the features contributing most strongly to Factor 1 across microbiome family composition (CLR-transformed) and circulating biomarkers (log10-transformed). Positive loadings (red) indicate positive associations with Factor 1, whereas negative loadings (blue) indicate inverse associations. Features are ordered within each view by absolute loading magnitude.

## LONG-TERM EXPERIMENT

### Body weights and fat

In the long-term experiment there were no significant differences in body weight increase between the *H. pylori* infected animals and control animals (**Fig. S2**), and there was no significant difference in perigonadal WAT weight between the two groups (males: 4.63%, 95% CI [−21.0, 39.3], females: −10.1%, 95% CI [−66.1, 178], posterior probabilities < 60%). The additional 17–18 weeks of HFD exposure, compared to the short-term experiment, likely resulted in control animals catching up to *H. pylori* animals in WAT weight. Overall, WAT weight increased approximately 8-fold in males and 2.5-fold in females over this period compared to the control animals in the short-term experiment.

### *H. pylori* infection is correlated with increased abundance of *Akkermansiaceae*

Akkermansiaceae expanded in the fecal microbiome of *H. pylori*–infected mice, particularly in males. At weaning, the relative abundance averaged 16.7% in males and 0.9% in females, compared to 0.04% in controls. Toward the end of the experiment, this increased to 31.96% in infected mice and 3.02% in non-infected controls (**Fig. S11**). Two of seven dams showed high Akkermansiaceae levels (56–62%), and pups from both of these dams were infected with *H. pylori* and had correspondingly higher Akkermansiaceae abundances (**Fig. S12**). We do not have data on the dam microbiome composition for the short-term experiment for comparison. Infected females showed higher CLR abundance of Clostridiaceae across time points and lower Lachnospiraceae five weeks post-diet change. Oscillospiraceae and Streptococcaceae were reduced at early time points in both sexes, while Tannerellaceae increased in the final fecal samples (**Fig. S11**). These patterns were consistent across colon, cecum, and stomach samples at termination **(Figs. S13–S15**). In the short-term experiment, gastric Helicobacteraceae abundance varied and was strongly associated with qPCR-estimated *H. pylori* load (slope 0.31, 95% CI [0.23, 0.40], ∼100%).

## DISCUSSION

In this mouse model, we show that early-life *H. pylori* infection accentuates the initial physiological response to HFD. This experimental finding complements our recent correlative study in Danish children and adolescents, in which *H. pylori* infection was associated with higher circulating glucose levels ^36^. During the short-term intervention, infection altered circulating metabolic biomarkers and, in the context of HFD, was associated with increased perigonadal adiposity and additional microbiome compositional changes, with the most pronounced effects observed in males. Notably, increased perigonadal WAT mass (**Fig. 2**) was not strongly reflected in overall body weight (**Fig. S2**), mirroring previous mouse studies in which early-life antibiotic-induced perturbations increased adiposity without weight gain ^4,72^ and aligning with human cross-sectional data ^73,74^. By contrast, the absence of differences in WAT mass following prolonged HFD exposure suggests that diet was the dominant determinant of adiposity at this stage.

Early-life *H. pylori* infection was associated with persistent alterations in endocrine responses in males. Elevated plasma leptin was the most consistent difference associated with *H. pylori* infection, and was detectable prior to the dietary switch (**Fig. 3**), as also reported in adult infection ^42^. The early elevation may reflect infection-induced regulation of leptin production. Infection can increase leptin secretion independently of adiposity ^75,76^, while *H. pylori* has additionally been shown to increase gastric leptin expression ^77,78^. Gastric leptin can also increase during HFD exposure and stimulate intestinal serotonin production, thereby promoting food intake ^79^. Leptin levels remained elevated in infected animals following the diet switch, particularly in males, where the difference persisted across all time points (**Fig. 4**).

In parallel, infected males showed higher ghrelin levels at the start of the diet intervention after a 4-hour fast. This finding is consistent with our recent observation that gastric organoids from infected animals show increased expression of endocrine cell markers, including ghrelin ^80^. Short-term fasting captures post-absorptive regulation and may reflect altered endocrine regulation during fasting, although the underlying mechanisms remain unclear. Insulin levels were also elevated in infected males receiving the HFD after two weeks (**Fig. 4**). However, leptin and ghrelin levels failed to respond as expected to a 12-hour fast (**Fig. 4**, **Fig. S3**). Together, these findings suggest altered endocrine responsiveness to fasting and disruption of normal leptin–ghrelin regulatory dynamics. The effect of fasting duration on ghrelin and leptin levels (4-hours vs 12-hours, **Fig. 4**) may contribute to discrepancies in associations with infection across studies.

The endocrine alterations were accompanied by context-dependent differences in inflammatory markers. The fact that at baseline, infected males had reduced MCP-1 levels together with borderline decreased TNF-α (**Fig. 3**), may represent a selective dampening of pro-inflammatory signaling, as a part of the tolerogenic phase of *H. pylori* infection that permits bacterial persistence while limiting excessive inflammation ^12^. The divergent metabolic and inflammatory responses (**Fig. 4, Fig. S3**) appear to indicate a shift toward inflammatory activation under metabolic stress. This transition may reflect the combined effects of adipose tissue expansion and sustained leptin elevation, which promote chemokine production and immune cell recruitment.

The high levels of MCP-1 after the 12-hour fast, the only biomarker values positively associated with *H. pylori* load **(Fig. S4**), are consistent with the known *H. pylori*-induction of MCP-1 secretion from gastric epithelial cells ^81–83^. In contrast, levels of IL-6 and other cytokines may reflect contributions from both infection-associated immune responses and adipose-derived inflammation ^84–86^. The concomitant rise in IL-10 at later stages could indicate attempted immune counter-regulation (**Fig. 4**), which co-occurred with inflammatory macrophage polarization in males receiving HFD (**Fig. S6B**). In females, the pattern differed: despite more limited adipose expansion (**Fig. 2**), infected animals showed elevated TNF-α under HFD (**Fig. 4**). Together, these findings highlight sex-specific responses to metabolic challenge, with males preferentially accumulating inflamed adipose tissue, whereas females exhibited a distinct pattern characterized by relatively limited adipose expansion together with elevated inflammatory markers. These findings are consistent with established sex differences in metabolic and immune regulation ^87^ that may be linked to reproductive fitness^88^.

*H. pylori* infection amplified HFD-induced shifts in intestinal microbiome composition. The resulting profile resembled a typical HFD-affected microbiome, with reduced fiber-degrading taxa and increased metabolically active, inflammation-associated bacteria ^89,90^. Lachnospiraceae, key short-chain fatty acid producers linked to energy harvest ^91,92^, were enriched, whereas Prevotellaceae, associated with fiber-rich diets and favorable metabolic profiles ^93–96^, were reduced. An increase in Streptococcaceae further suggests a disrupted or inflammatory microbial niche ^97,98^ and has previously been linked to *H. pylori* infection ^99–101^. Overall, these findings indicate that microbiome shifts are primarily diet-driven, suggesting that the influence of *H. pylori* on intestinal microbiome composition is context dependent and occurs against a strong dietary background. This is supported by the expansion of Akkermansiaceae in infected animals only in the long-term experiment (**Fig. S10B and 11B**). Differences in relative abundances were apparent prior to the diet switch and likely maternally transferred (**Fig. S12**), and the expansion may reflect opportunistic growth under diet-altered conditions. The role of *Akkermansia* in the microbiome and on host physiology is complex; indeed, *Akkermansia* has been shown to act opportunistically when the gut microbiome is disrupted ^1,102,103^.

Exposure to high-fat, calorie-dense diets represents a novel challenge within the evolutionary timescale of the *H. pylori*–host relationship. For most of human evolutionary history, nutritional environments were likely characterized by fluctuating cycles of abundance and scarcity ^104^. In such contexts, infection-associated metabolic phenotypes favoring efficient energy storage during periods of abundance may have conferred a selective advantage ^105^, potentially contributing to the long-term evolutionary persistence and global ubiquity of the human–*H. pylori* association.

Several limitations point toward important directions for future work. Our data identify coordinated endocrine, inflammatory, adipose, and microbiome alterations associated with early-life *H. pylori* infection but do not establish the causal relationships among these processes. Determining how infection-induced endocrine signaling interacts with immune responses, the intestinal microbiome, and sex-specific regulatory pathways to influence energy allocation and adipose expansion will be essential for understanding the observed phenotype. In addition, because circulating hormones and cytokines were not measured in the long-term experiment, it remains unclear whether the endocrine alterations observed during the initial response to HFD persist despite the convergence in adiposity. Future studies combining longitudinal metabolic phenotyping with mechanistic manipulation of endocrine, immune, and microbial pathways will be important for defining how early-life *H. pylori* infection modifies host responses to dietary challenge.

## Data availability

The datasets supporting the conclusions of this article are available in the Sequence Read Archive (SRA) repository, under the Bioproject ID PRJNA1207418 (https://www.ncbi.nlm.nih.gov/bioproject/PRJNA1207418) and Zenodo repository (https://doi.org/10.5281/zenodo.19671081). The code is available at GitHub (https://github.com/sbreum/Early-life-Helicobacter-pylori-infection-worsens-metabolic-state-in-mice-receiving-a-high-fat-diet). The work in this manuscript is available as a preprint, and has been presented as a poster at a conference ^106,107^.

## Acknowledgements

We thank Danai-Anastasia Panou for valuable feedback on the final draft. We also extend our gratitude to the following facilities at the University of Copenhagen for their expertise, support and infrastructure: the animal caretakers at the 16.2 unit, Department for Experimental Medicine; the staff at GeoGenetics Sequencing Core, Globe Institute; Tina Brand, Pernille Selmer Olsen and Lasse Vinner at the Molecular Biology Labs, Globe Institute; and the staff at FACS Core Facility, Faculty of Health and Medical Sciences, University of Copenhagen. We thank Professor Anne Müller at the Institute for Molecular Cancer Research, University of Zurich for assistance with establishing the neonatal mouse infection model. We used BioRender with license Andersen, S. (2025) to create figure 1 (https://BioRender.com/ngo64c1) and 2 (https://BioRender.com/14tvdxy). ChatGPT (GPT- 5.5) was used for copyediting and language refinement, enhancing idea generation and exploration of metabolic connections, and evaluation of analytical code.

