## Supplementary Figs+Tables+Methods for "Early-life *Helicobacter pylori* infection in mice alters endocrine signaling and accelerates adipose expansion during high-fat feeding"

**SUPPLEMENTARY MATERIALS**

**SUPPLEMENTARY FIGURES**

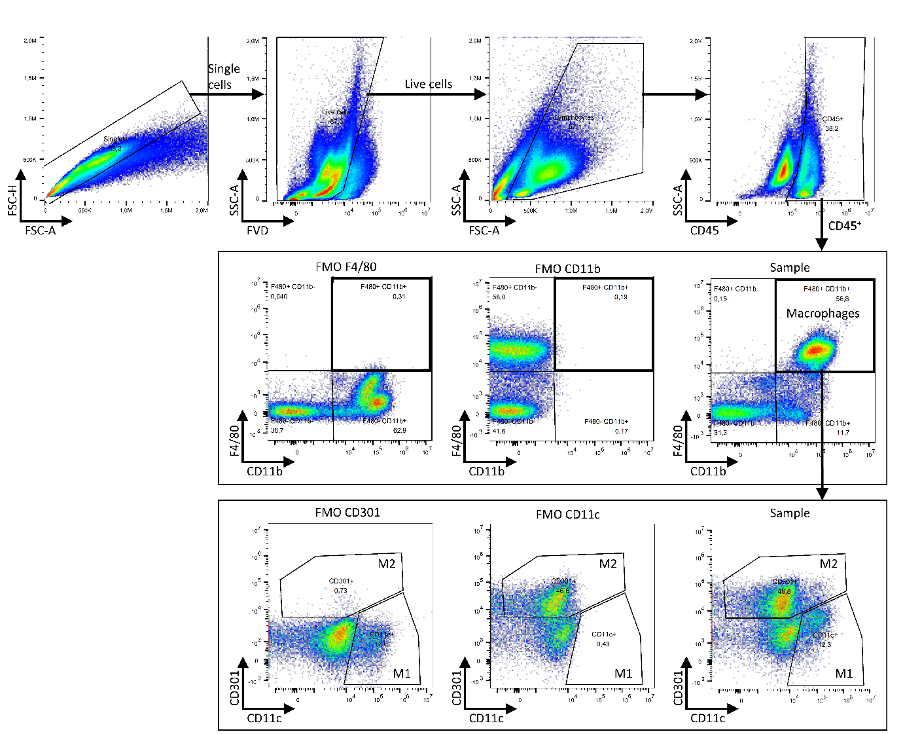

**Figure S1:** **Short-term experiment**. Flow cytometry gating strategy (example from control mouse). Cells from the stromal vascular fraction isolated from epididymal white adipose tissue depots (right depot from males and both depots from females) were first gated for single cells (FSC-H vs. FSC-A). Singlets were further gated for viable cells based on staining with a fixable viability dye (FVD). Cell debris was excluded based on SSC-A vs. FSC-A pattern and leukocytes were gated based on CD45 expression. From this population, macrophages were gated based on surface expression of F4/80 and CD11b. Gates were set based on Fluorescence Minus One (FMO) controls. F4/80^+^ CD11b^+^ macrophages were finally analyzed for CD301 and CD11c surface expression to characterize “pro-inflammatory” M1 (CD11c^+^) and “anti-inflammatory” M2 (CD301^+^) macrophage subpopulations based on FMO controls.

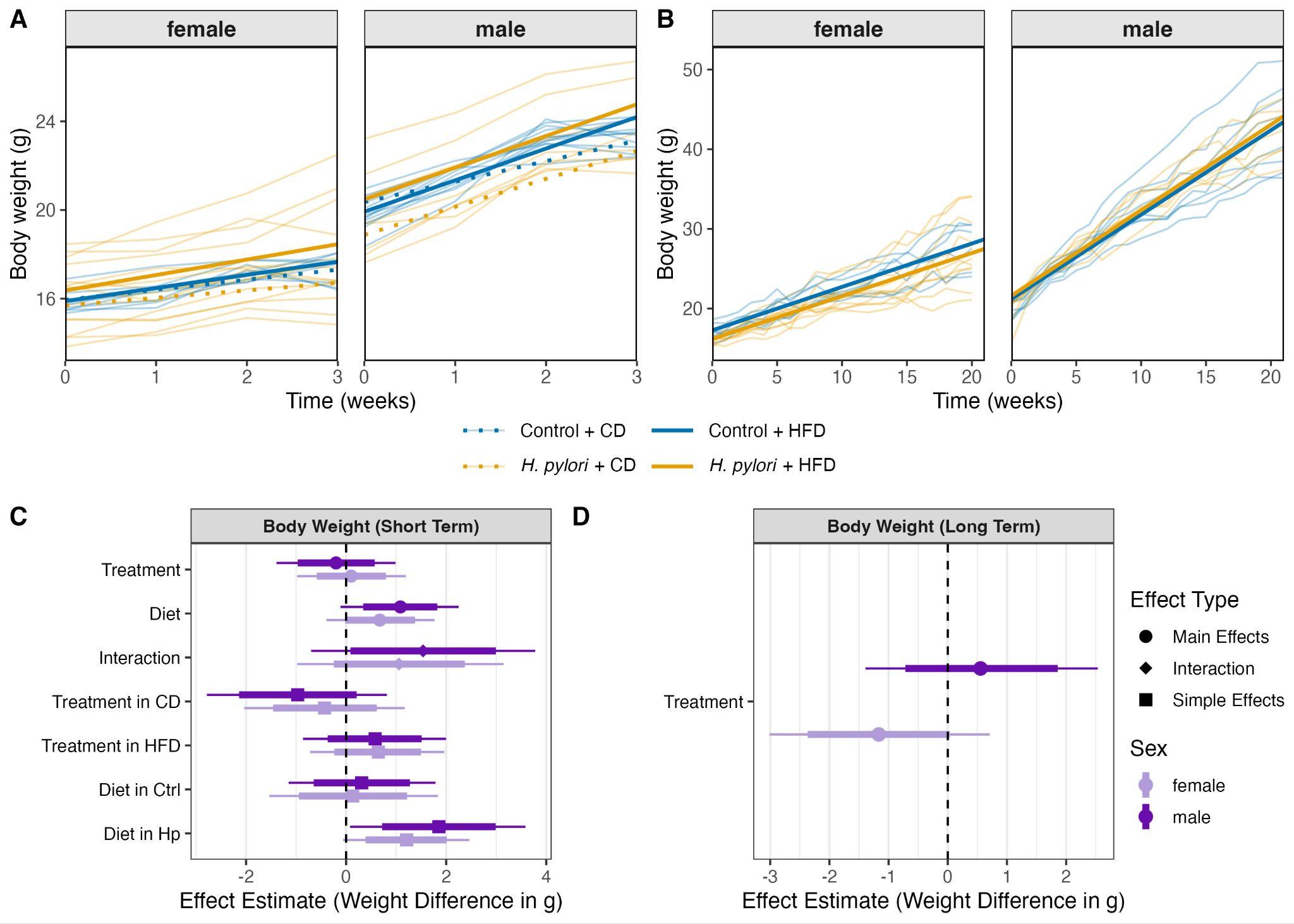

**Figure S2:** Effect of early-life *H. pylori* infection on relative body weight gain in C57BL/6JRj mice. **A** Time trajectory of relative body weight gain in a short-term experiment of mice receiving a control diet (CD) or high-fat diet (HFD) for three weeks. The bold lines represent the average body weight gain for control mice on a CD (blue dotted line, n = 3 females and 4 males) or on a HFD (blue solid, n = 5 females and 6 males), or for *H. pylori* infected mice on a CD (yellow dotted, n = 6 females and 3 males) or on a HFD (yellow solid, n = 8 females and 5 males). **B** Time trajectory of relative body weight gain in the long-term experiment of mice for 20 weeks (females) and 21 weeks (males) on HFD. Bold lines represent the average body weight gain for *H. pylori* infected mice (yellow, n = 11 females and 7 males) and control mice (blue, n = 6 females and 9 males). All received a HFD. **C** Forest plot of the results from a Bayesian mixed-effects regression with time (weeks) as a continuous predictor. In the short-term experiment, when comparing the male mice infected with *H. pylori*, animals receiving a HFD were heavier than mice receiving the control diet. **D** Treatment did not result in a weight difference in the long-term experiment. Forest plots showing posterior median effect estimates for main effects (treatment, diet, and their interaction; *H. pylori* vs control) and corresponding simple effects (treatment within each diet and diet within each treatment). Error bars show 95% credible intervals (thin lines) and 80% credible intervals (thick lines). Effect types are indicated by symbol shape (dot, main effects; diamond, interaction effects; square, simple effects), while light and dark purple denote females and males, respectively. The horizontal dashed line indicates no effect.

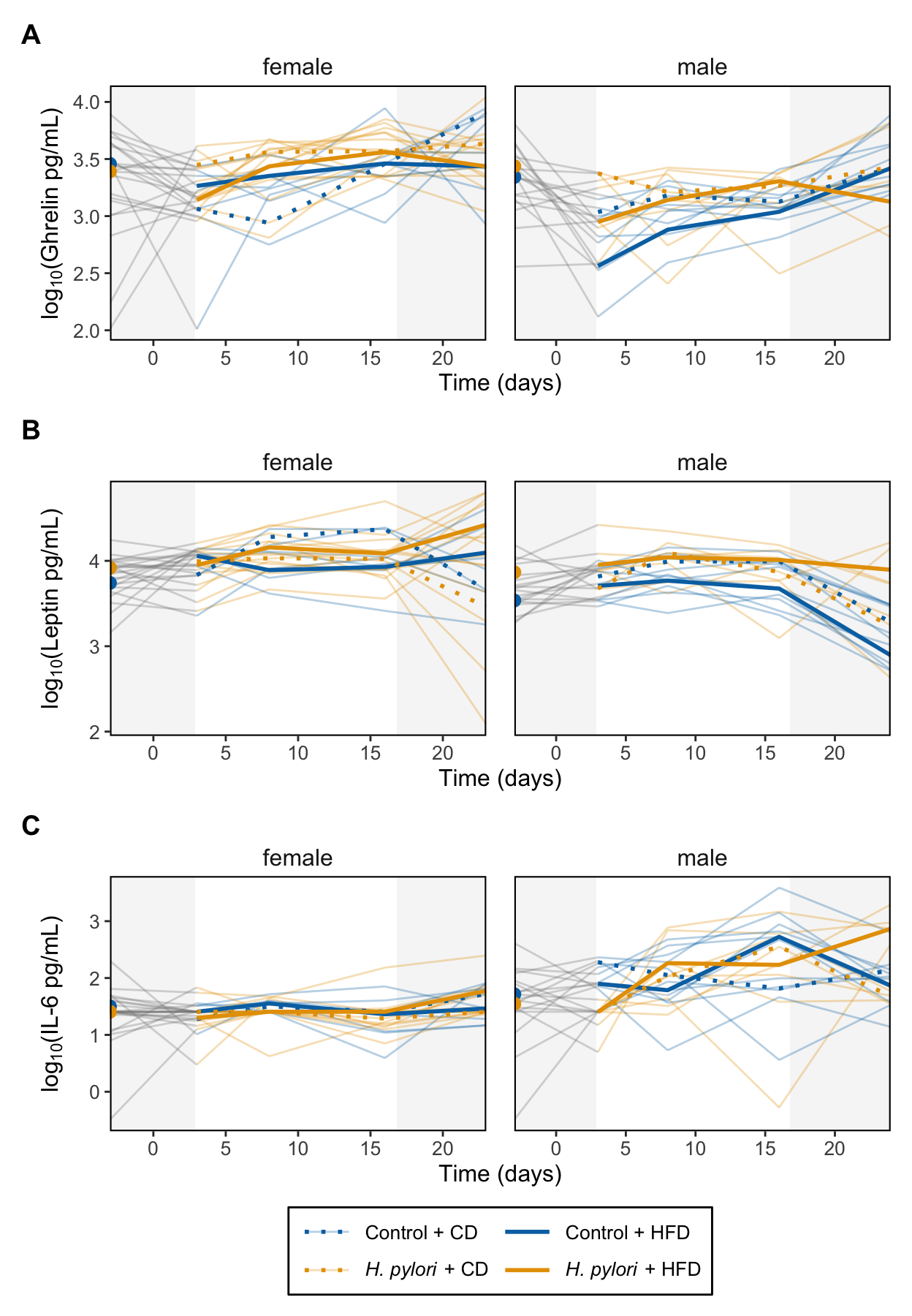

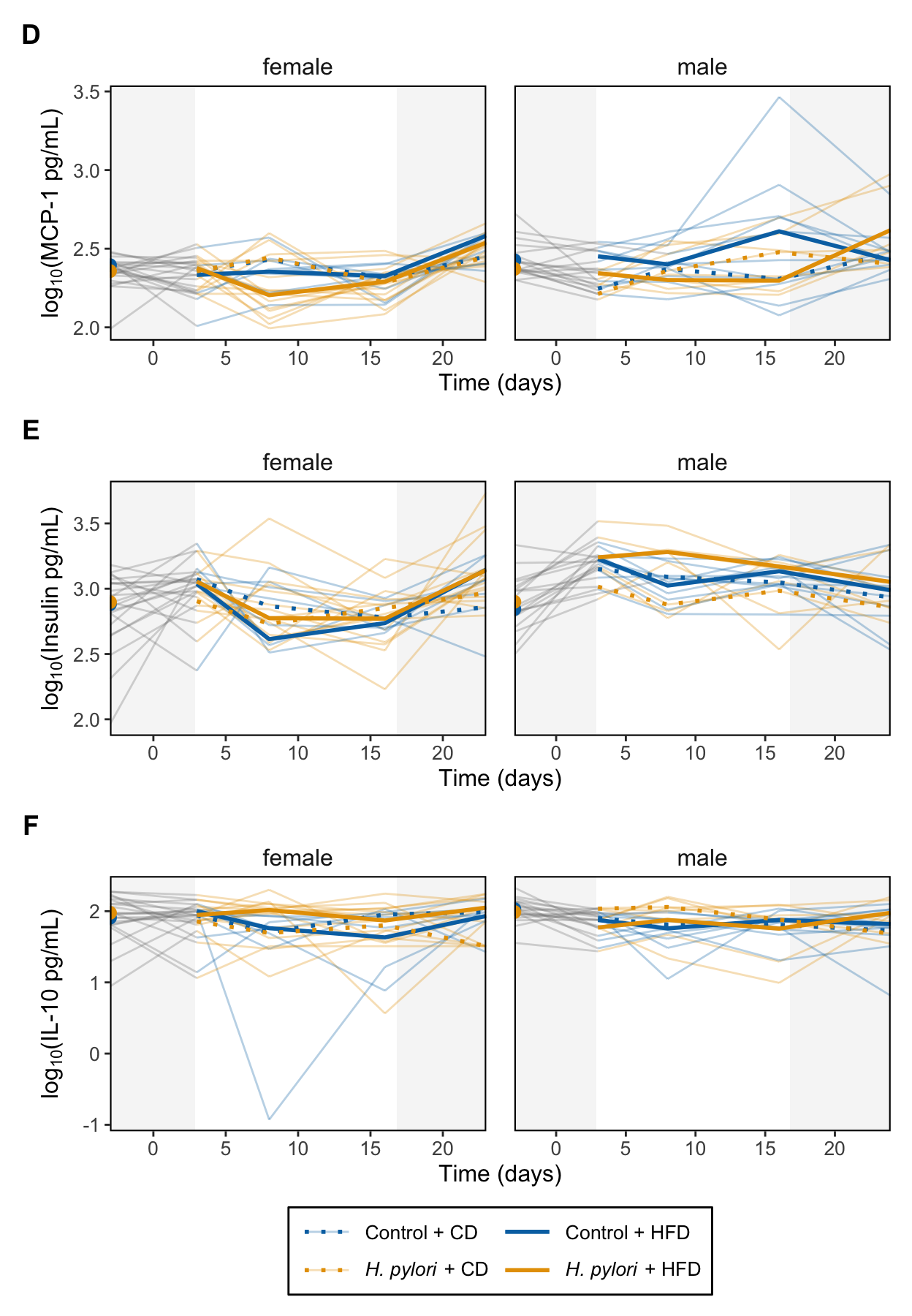

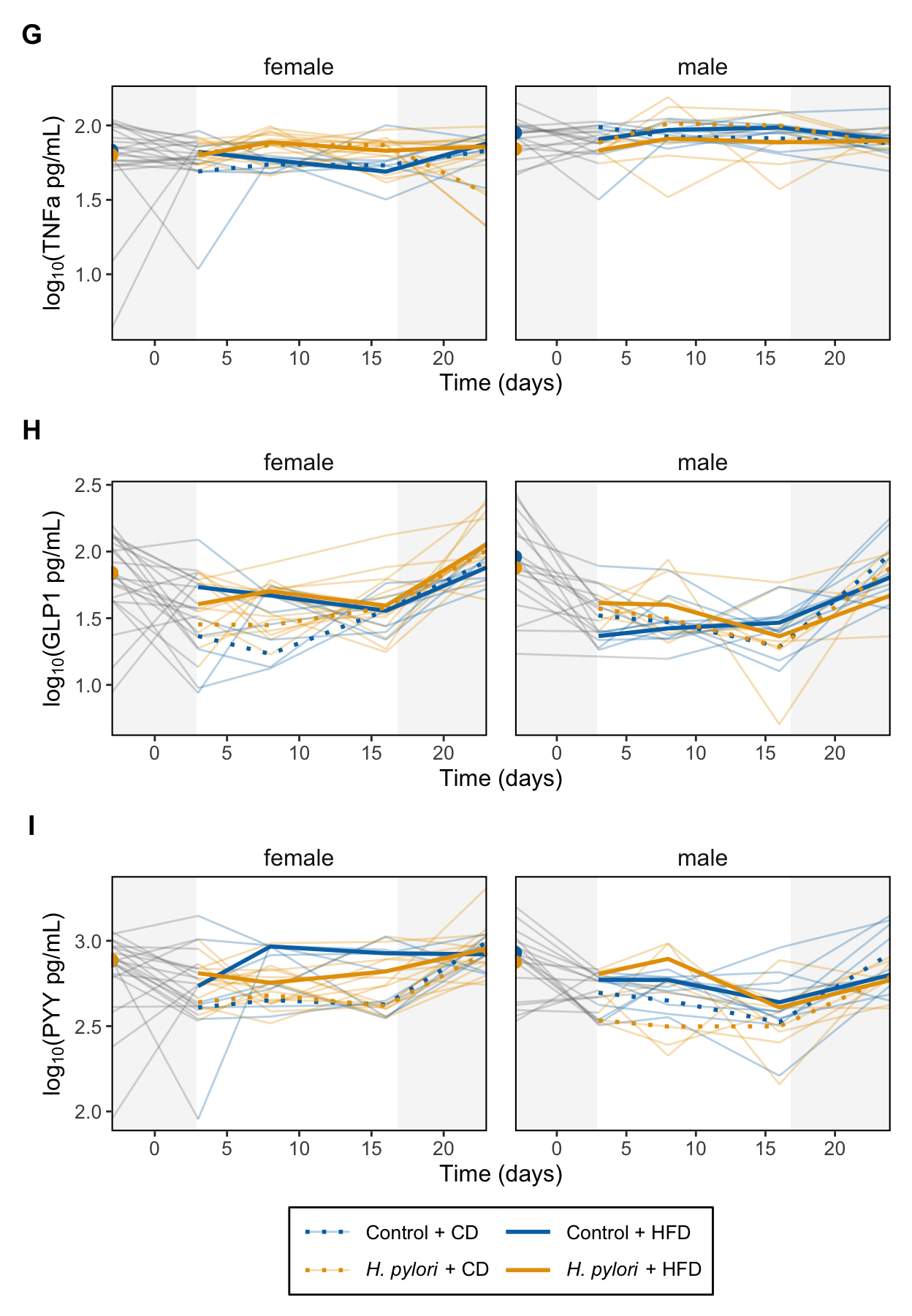

**Figure S3: Short-term experiment.** Effect of early-life *H. pylori* infection on circulating **A** ghrelin, **B** leptin, **C** IL-6, **D** MCP-1, **E** insulin, **F** TNF-α, **G** PYY, **H** IL-10 and **I** GLP-1 before and after high-fat diet (HFD) intervention in C57BL/6JRj mice. On the x-axis is time in days, and biomarker levels are shown as log10-transformed concentrations. Grey blocks mark samples collected 3 days before the diet intervention, and on day 24 at termination of the experiment. The first four samples were collected after a 4h fast, and at termination after a 12h fast. The bold lines represent the median biomarker concentration for control mice on a CD (blue dotted line, n = 3 females and 4 males) or on a HFD (blue dashed, n = 5 females and 6 males), or for *H. pylori* infected mice on a CD (yellow dotted, n = 6 females and 3 males) or on a HFD (yellow dashed, n = 8 females and 5 males).

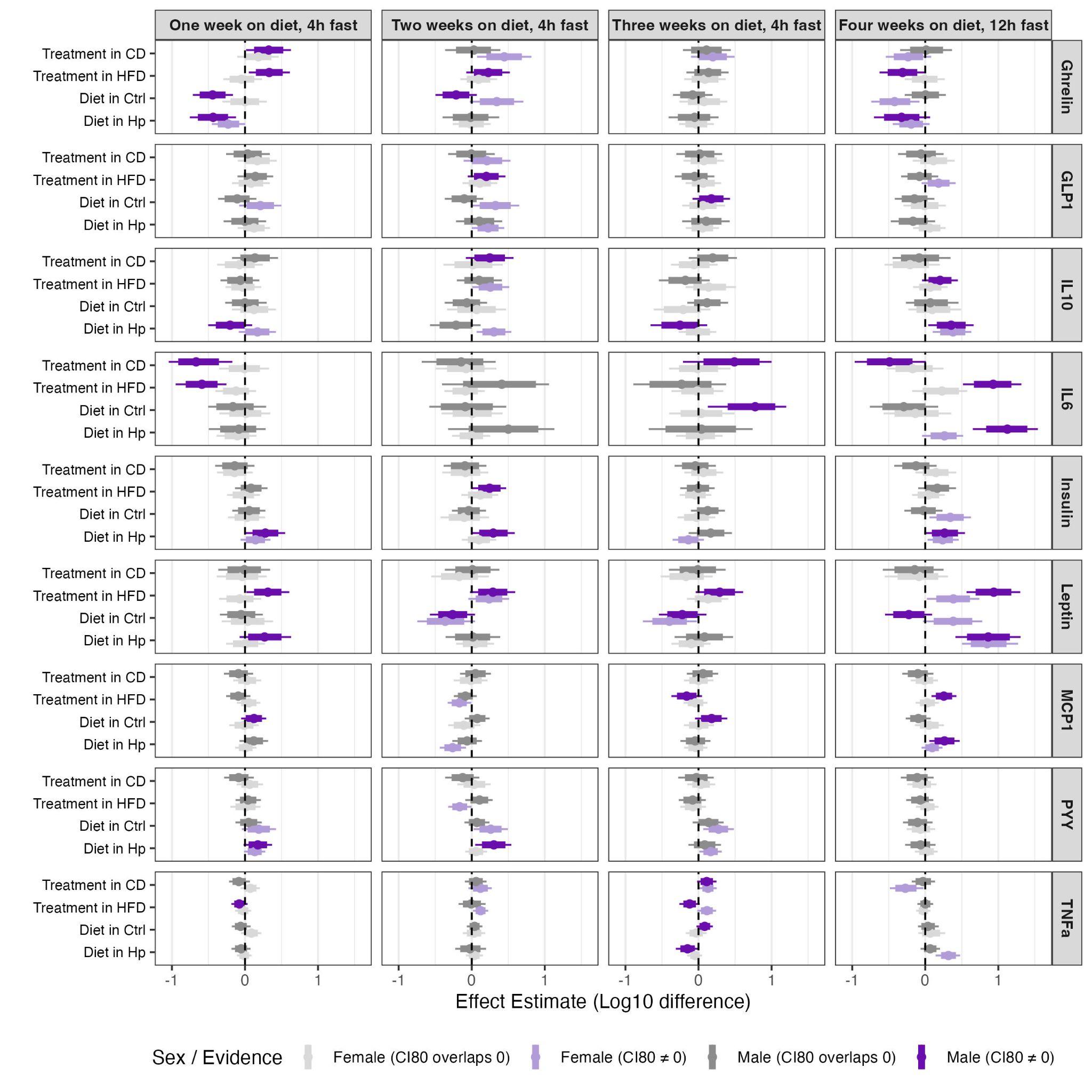

**Figure S4:** **Short-term experiment.** Posterior estimates of simple effects for nine biomarkers across four sampling timepoints between *H. pylori*–infected and control animals sampled one to four weeks after switching to a high-fat diet (HFD) or control diet. Points show posterior median effect estimates and error bars show 95% credible intervals (thin lines) and 80% credible intervals (thick lines). Effects are expressed as posterior median differences in biomarker concentration in log_10_ units. Simple effects shown are treatment within diet (Hp − Ctrl in CD and HFD) and diet within treatment (HFD − CD in Ctrl and Hp), derived from the full Bayesian mixed-effects model including treatment, diet, sex, and categorical timepoint interactions, with a random intercept for mouseID to account for repeated measurements. Panels are arranged by biomarker (rows) and timepoint (columns). For samples collected one to three weeks post diet switch, animals were fasted for 4 h; for the final timepoint, animals were fasted for 12 h. Light and dark purple indicate female and male estimates, respectively, where the 80% credible interval excluded zero; light and dark grey indicate female and male estimates, respectively, where the 80% credible interval overlapped zero. The horizontal dashed line indicates no effect.

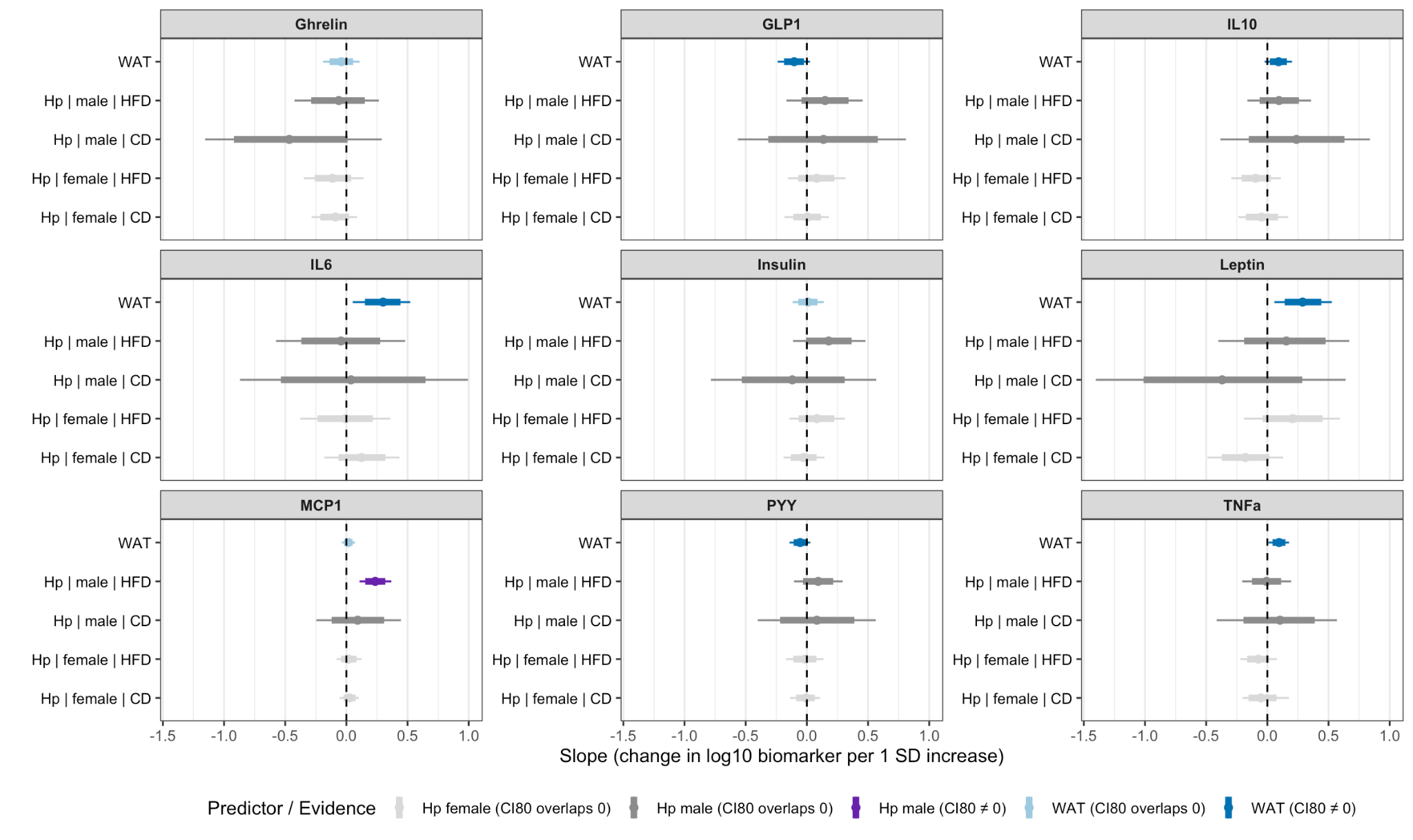

**Figure S5**: **Short-term experiment.** Association between white adipose tissue (WAT) weight and *H. pylori* load of C57BL/6JRj mice across nine biomarkers. Results are based on a Bayesian model testing associations between log_10_-transformed biomarker concentrations measured at termination following a 12 h fast, standardized WAT weight (g), and log_10_-transformed *H. pylori* load quantified by qPCR of gastric tissue. Forest plots display posterior median effect estimates (points) with 95% credible intervals (thin lines) and 80% credible intervals (thick lines). For *H. pylori* load, light and dark purple indicate female and male estimates, respectively, where the 80% credible interval excludes zero, while light and dark grey indicate estimates where the 80% credible interval overlaps zero. For WAT weight, dark blue indicates estimates where the 80% credible interval excludes zero and light blue where it overlaps zero. The horizontal dashed line denotes no effect.

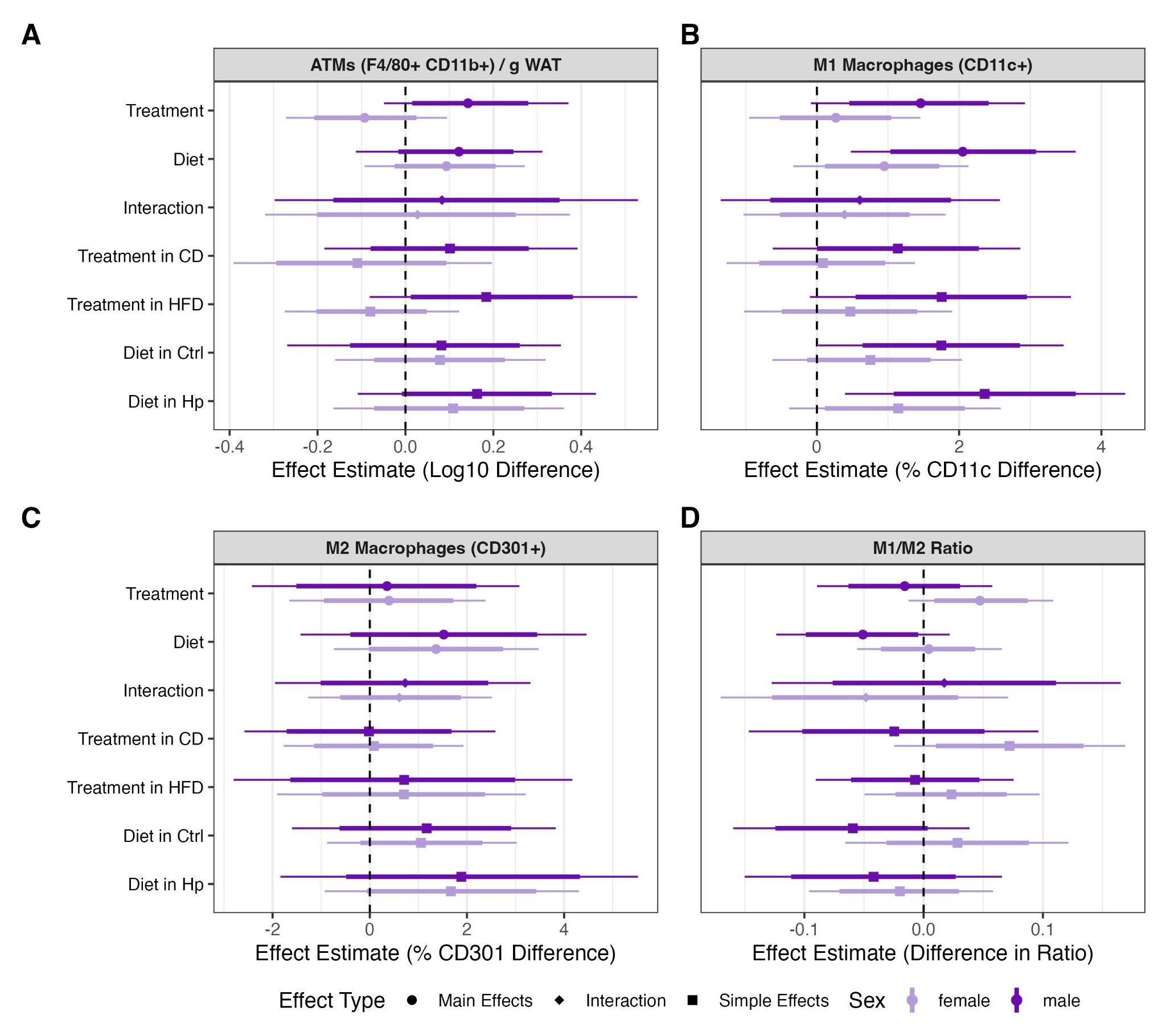

**Figure S6**: **Short-term experiment.** Effect of early-life *H. pylori* infection on macrophage subtypes in perigonadal white adipose tissue (WAT) of C57BL/6JRj mice three weeks after receiving a control or high-fat diet. Forest plots of the results of Bayesian regression models testing the effects of treatment (*H. pylori* infection or control) and diet (HFD or CD) on **A** log_10_-transformed counts of F4/80^+^ CD11b^+^ macrophages (adipose tissue macrophages) per gram perigonadal WAT. **B** Frequencies of CD11c^+^ macrophages out of F4/80^+^ CD11b^+^ macrophages. **C** Frequencies of CD301^+^ macrophages as a proportion of F4/80^+^ CD11b^+^ macrophages. **D** Ratio of CD11c^+^ to CD301^+^ macrophages. Effect types are indicated by symbol shape (dot, main effects; diamond, interaction effects; square, simple effects), with 95% credible intervals (thin lines) and 80% credible intervals (thick lines), while light and dark purple denote females and males, respectively.

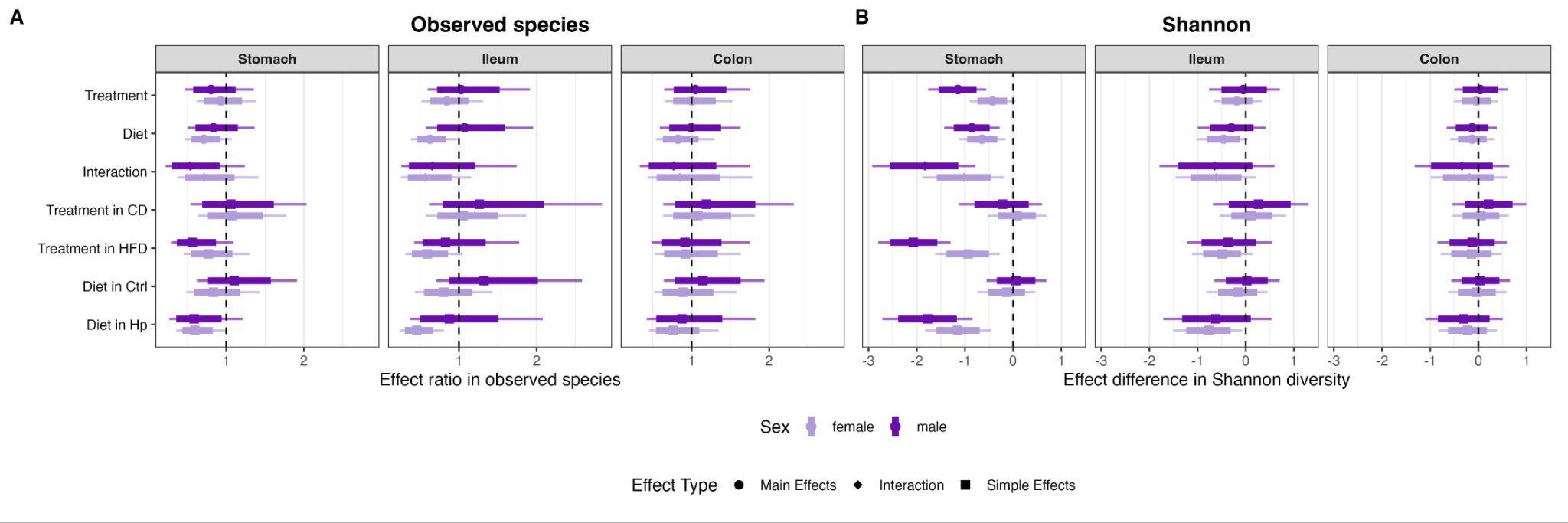

**Figure S7: Short-term experiment.** Effects of early-life *H. pylori* infection on gastrointestinal alpha diversity in C57BL/6JRj mice. Forest plots of the effects of treatment (control versus *H. pylori*) and diet (control diet (CD) or high fat diet (HFD)) on the alpha diversity. Treatment and diet interacted to lower Shannon diversity of the stomach microbiomes of *H. pylori* males and females receiving a HFD. **A** Observed Species and **B** Shannon Diversity Index of stomach, ileum and colon microbiomes three weeks after change to HFD. Forest plots showing posterior median effect estimates for main effects (treatment, diet, and their interaction; *H. pylori* vs control) and corresponding simple effects (treatment within each diet and diet within each treatment). Error bars show 95% credible intervals (thin lines) and 80% credible intervals (thick lines). Effect types are indicated by symbol shape (dot, main effects; diamond, interaction effects; square, simple effects), while light and dark purple denote females and males, respectively. The horizontal dashed line indicates no effect.

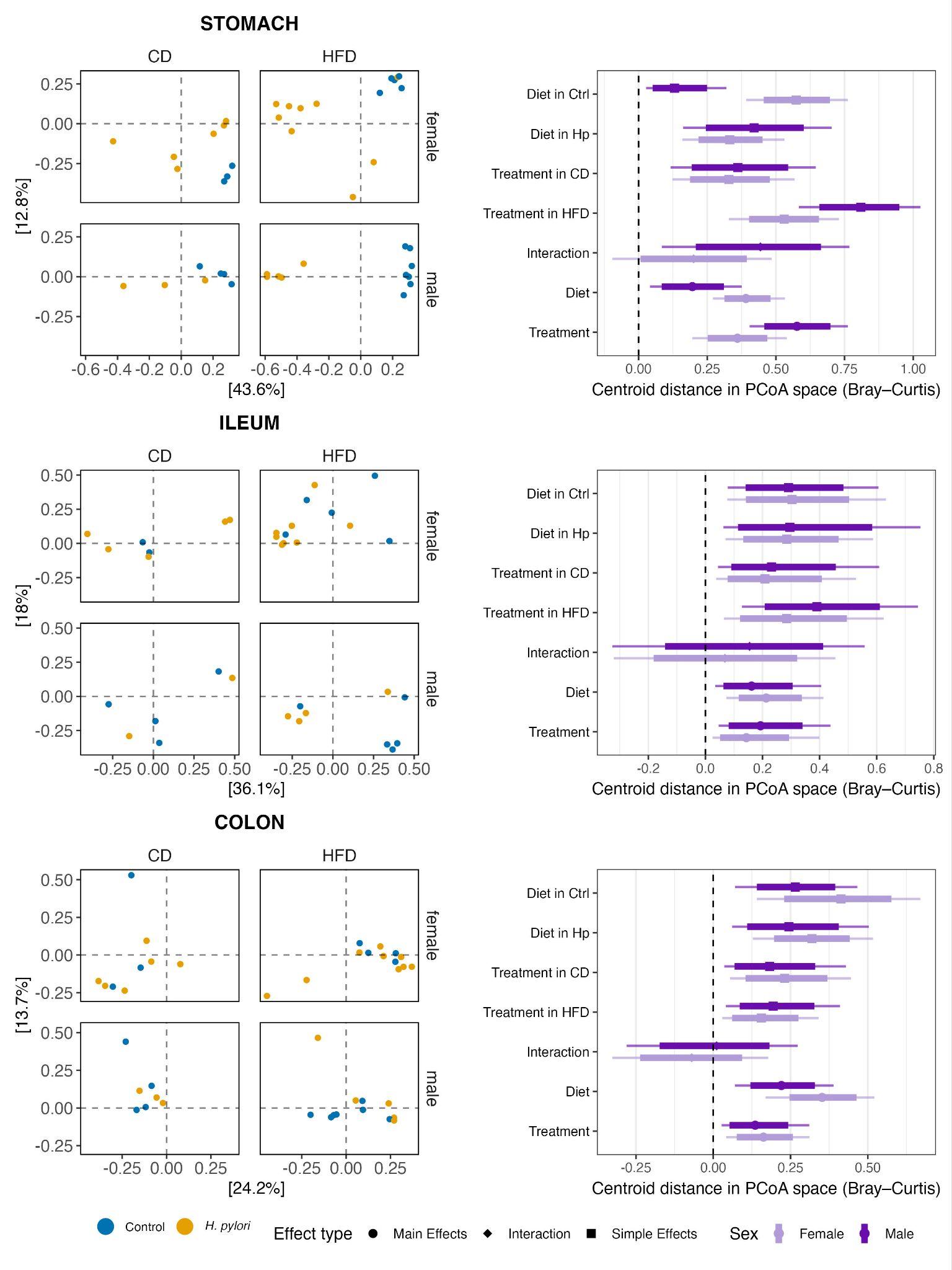

**Figure S8: Short-term experiment.** **Early-life *H. pylori* infection primarily affects gastric microbiome composition.** Principal coordinate analysis (PCoA) of Bray–Curtis dissimilarity based on relative abundances illustrates differences in microbial community composition among samples collected from the stomach, ileum, and colon at the end of the experiment, three weeks after transition to control diet (CD) or high-fat diet (HFD). Samples are stratified by sex and diet, and colored by infection status. For each gastrointestinal site, corresponding forest plots summarize Bayesian model–derived estimates of community separation in PCoA space, shown as posterior median centroid distances with 95% credible intervals. These estimates quantify the degree of multivariate community separation rather than effect sizes in the classical regression sense. Diet and *H. pylori* infection were associated with increased separation of microbial communities in ordination space. The reported estimates reflect the magnitude of community dissimilarity rather than changes in individual taxa or absolute effect sizes. Effect types are marked by shape with a dot for main effects, a diamond for interactions and a square for simple effects, while light and dark purple denotes females and males. Panels show stomach (A–B), ileum (C–D), and colon (E–F).

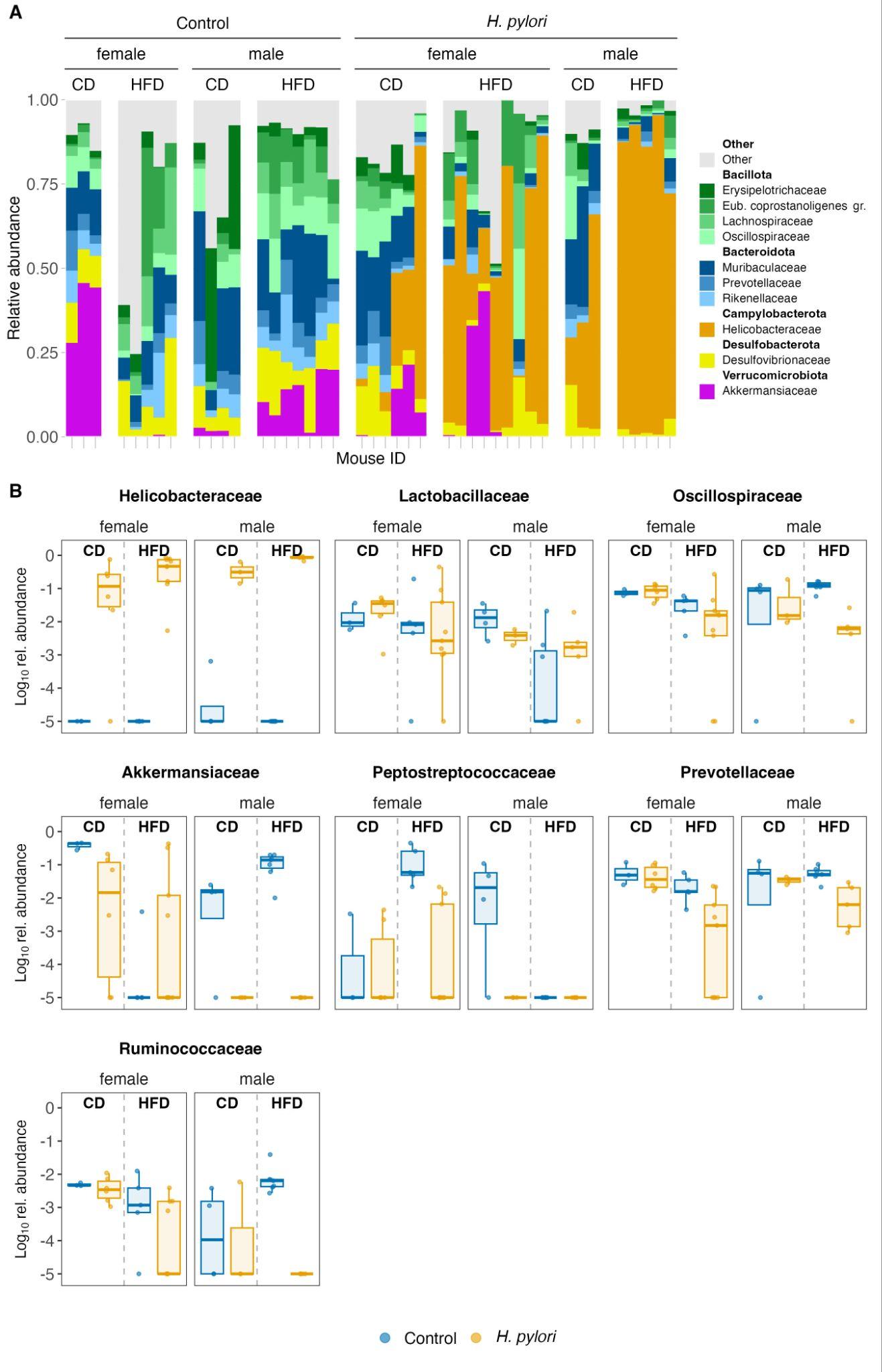

**Figure S9: Short-term experiment.** Gastric microbiome composition in *H. pylori* infected or control C57BL/6JRj mice, measured by 16S rRNA gene sequencing. The experiment ended three weeks after a change to control diet (CD) or high-fat diet (HFD). **A** Bars represent stacks of the 10 most relative abundant bacterial families in individual mice and are sorted in panels according to treatment, sex and diet. The colors indicate the phylum, and the gradient of a given color represents different families within the phylum. **B** The nine taxonomic gastric families among the top 10 most abundant, which showed differences between *H. pylori*-infected (yellow) and control (blue) mice, stratified by sex and diet. The relative abundances were log_10_-transformed prior to the analysis.

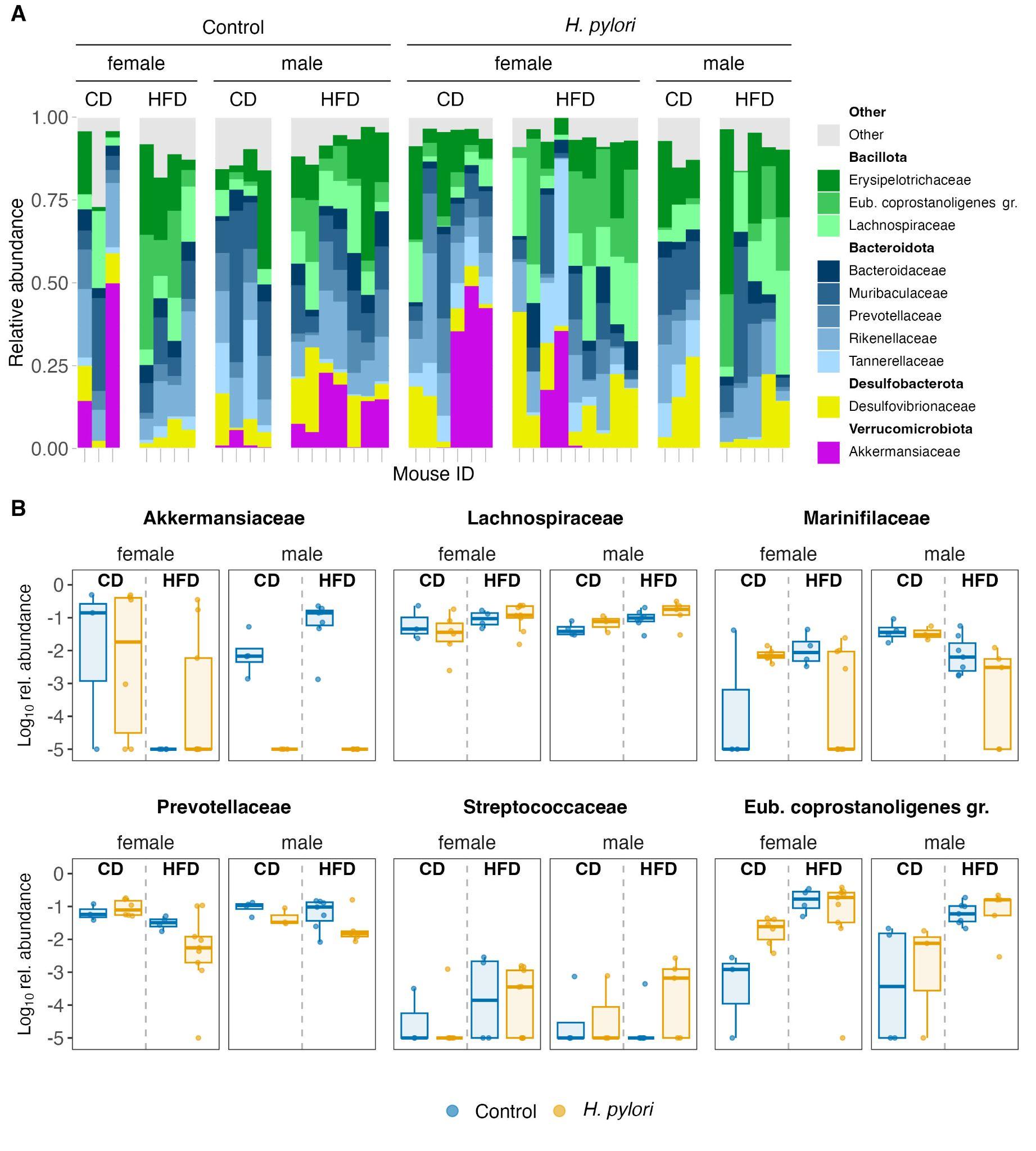

**Figure S10:** **Short-term experiment.** Colonic microbiome composition in *H. pylori* infected or control C57BL/6JRj mice, measured by 16S rRNA gene sequencing. The experiment ended three weeks after a change to control diet (CD) or high-fat diet (HFD). **A** Bars represent stacks of the 10 most relatively abundant bacterial families in individual mice and are sorted in panels according to treatment, sex and diet. The colors indicate the phylum and the gradient of a given color represents different families within the phylum. **B** Six colonic families, which showed differences in prevalence between *H. pylori*-infected (yellow) and control (blue) mice, stratified by sex and diet. The relative abundances were log_10_-transformed prior to the analysis.

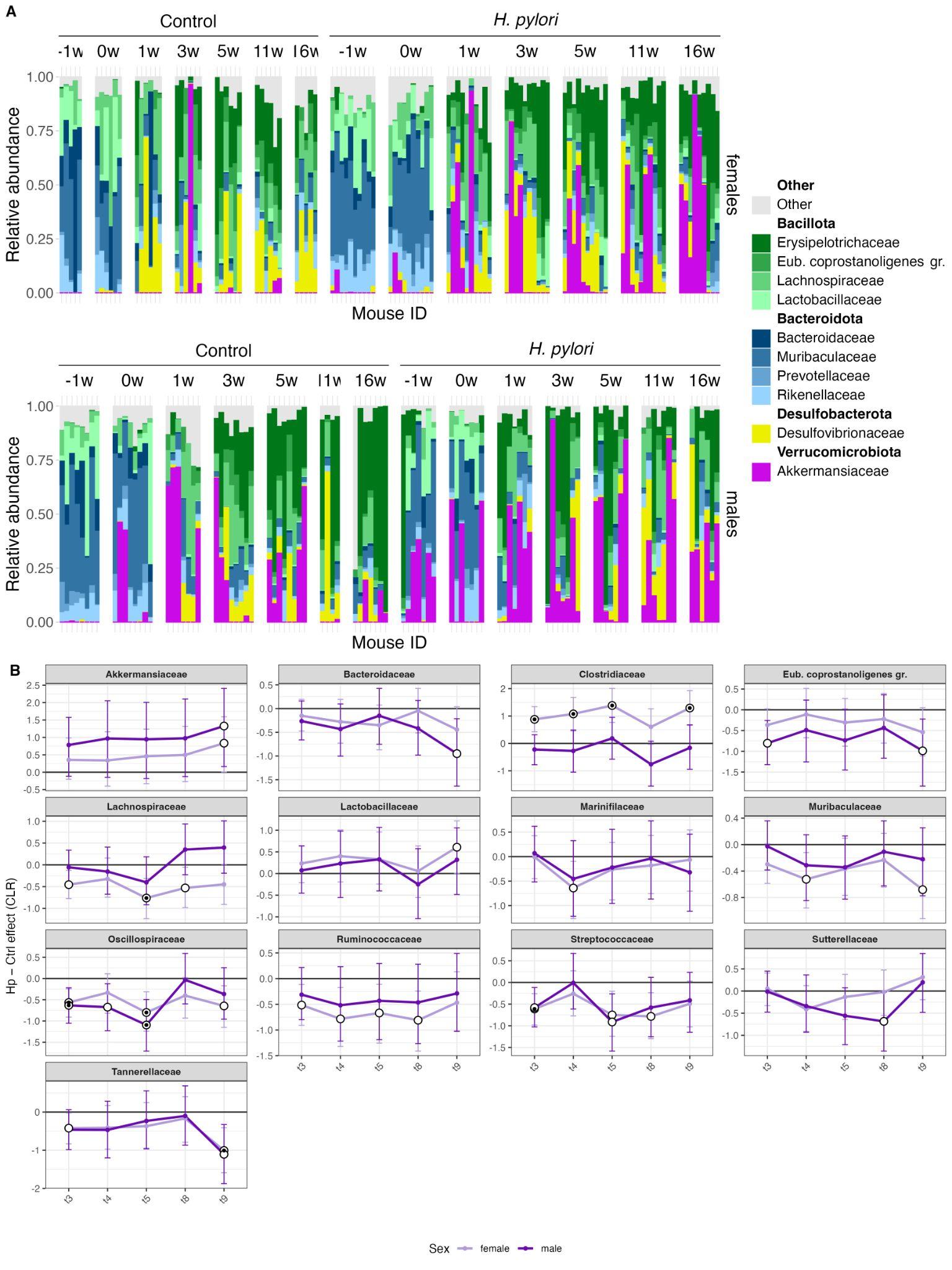

**Figure S11: Long-term experiment.** Early-life *Helicobacter pylori* infection correlates with changes in the composition of the fecal microbiome composition compared to C57BL/6JRj control mice. **A** Long-term experiment time trajectory of fecal microbiome composition in *H. pylori* infected and control C57BL/6JRj mice by sex, measured by 16S rRNA gene sequencing. Bars represent the 10 most abundant bacterial families in panels representing mice grouped per time point relative to diet change from chow to high-fat diet. Panels are sorted after treatment (Control vs. *H. pylori*). Bacterial families belonging to the same phylum are shown as a variation of similar colors. W = week. **B** Trajectories of differences in CLR-transformed abundances of bacterial families between *H. pylori*–infected and control animals sampled three to nine weeks after switching to a high-fat diet. The y-axis shows the effect size as the posterior median difference between treatments, expressed in centered log-ratio (CLR) units. These represent the simple effects of treatment within each sex, estimated from a Bayesian mixed-effects model including treatment, sex, and timepoint interactions, with a random intercept for mouseID to account for repeated measurements. Thus, an effect estimate of 0.5 corresponds to approximately a 3.2-fold change in relative abundance (on the CLR scale) in *H. pylori*–infected animals compared with controls. Because values represent differences between groups, a positive effect may reflect either an increase in infected animals or a decrease in controls. Error bars represent 95% credible intervals. White circles indicate estimates where the 80% credible interval excludes zero, and black dots indicate estimates where the 95% credible interval excludes zero. Only families where at least one sample had a 80% credible interval that did not cross 0 are shown. Light and dark purple denote females and males, respectively.

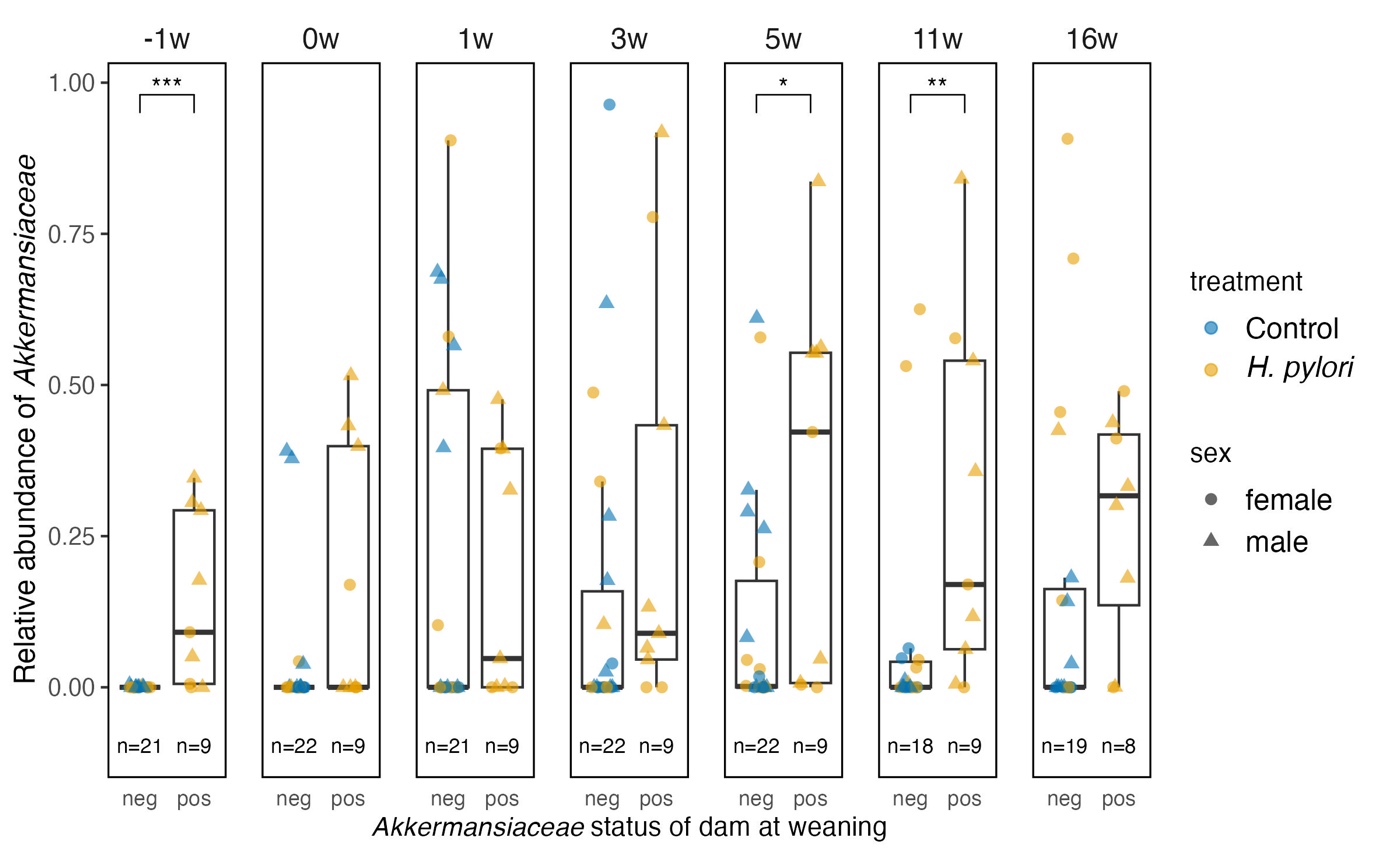

**Figure S12: Long-term experiment.** Effect of Akkermansiaceae status of C57BL/6JRj dam at weaning, on the relative abundance of Akkermansiaceae in the feces of the weaned pups over time. Of seven dams, two were Akkermansiaceae positive. Statistical differences were determined by Wilcoxon rank-sum test: *: p<0.05, **: p<0.01, ***: p<0.001. Abbreviations: w; weeks, neg; negative, pos; positive

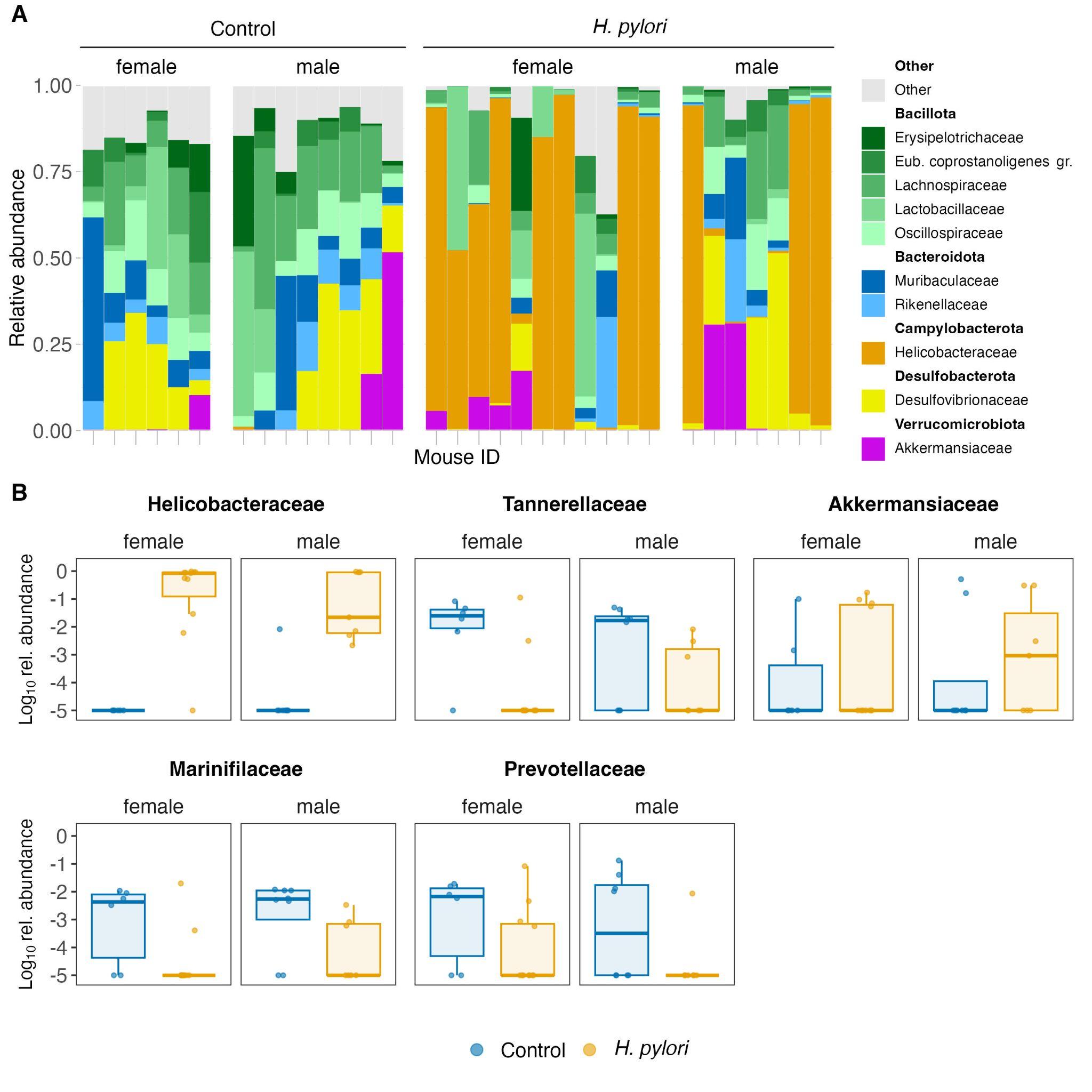

**Figure S13: Long-term experiment.** Stomach microbiome composition in *H. pylori* infected or control C57BL/6JRj mice, measured by 16S rRNA gene sequencing. **A** Bars represent stacks of the 10 most relative abundant bacterial families in individual mice and are sorted in panels according to sex and treatment. The colors indicate the phylum, and the gradient of a given color represents different families within the phylum. **B** Nine bacterial families, which showed differences in prevalence between *H. pylori*-infected (yellow) and control (blue) mice, stratified by sex. The relative abundances were log_10_-transformed prior to the analysis.

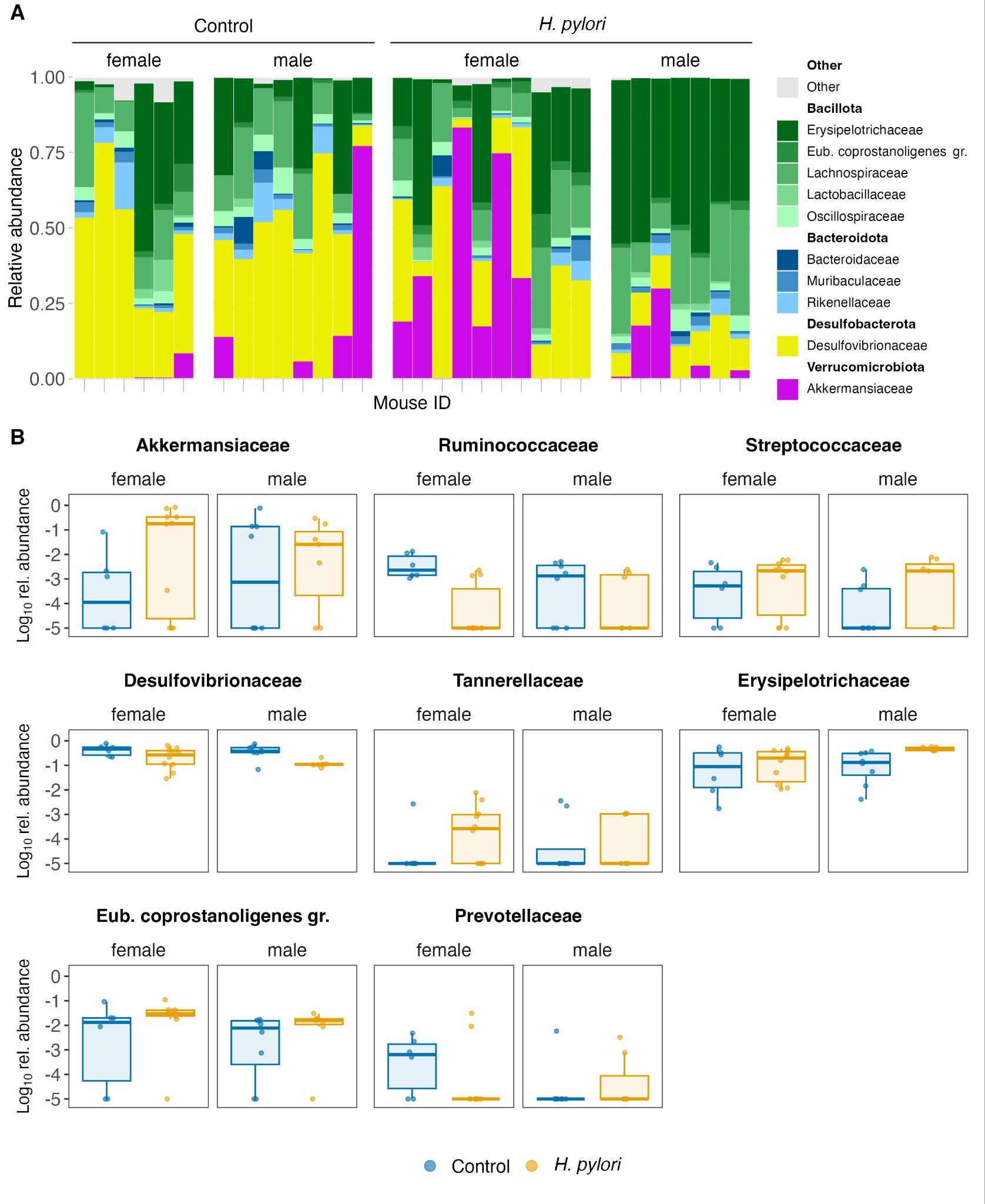

**Figure S14: Long-term experiment.** Cecum microbiome composition in *H. pylori* infected or control C57BL/6JRj mice, measured by 16S rRNA gene sequencing. **A** Bars represent stacks of the 10 most relative abundant bacterial families in individual mice and are sorted in panels according to sex and treatment. The colors indicate the phylum, and the gradient of a given color represents different families within the phylum. **B** Nine bacterial families, which showed differences in prevalence between *H. pylori*-infected (yellow) and control (blue) mice, stratified by sex. The relative abundances were log_10_-transformed prior to the analysis.

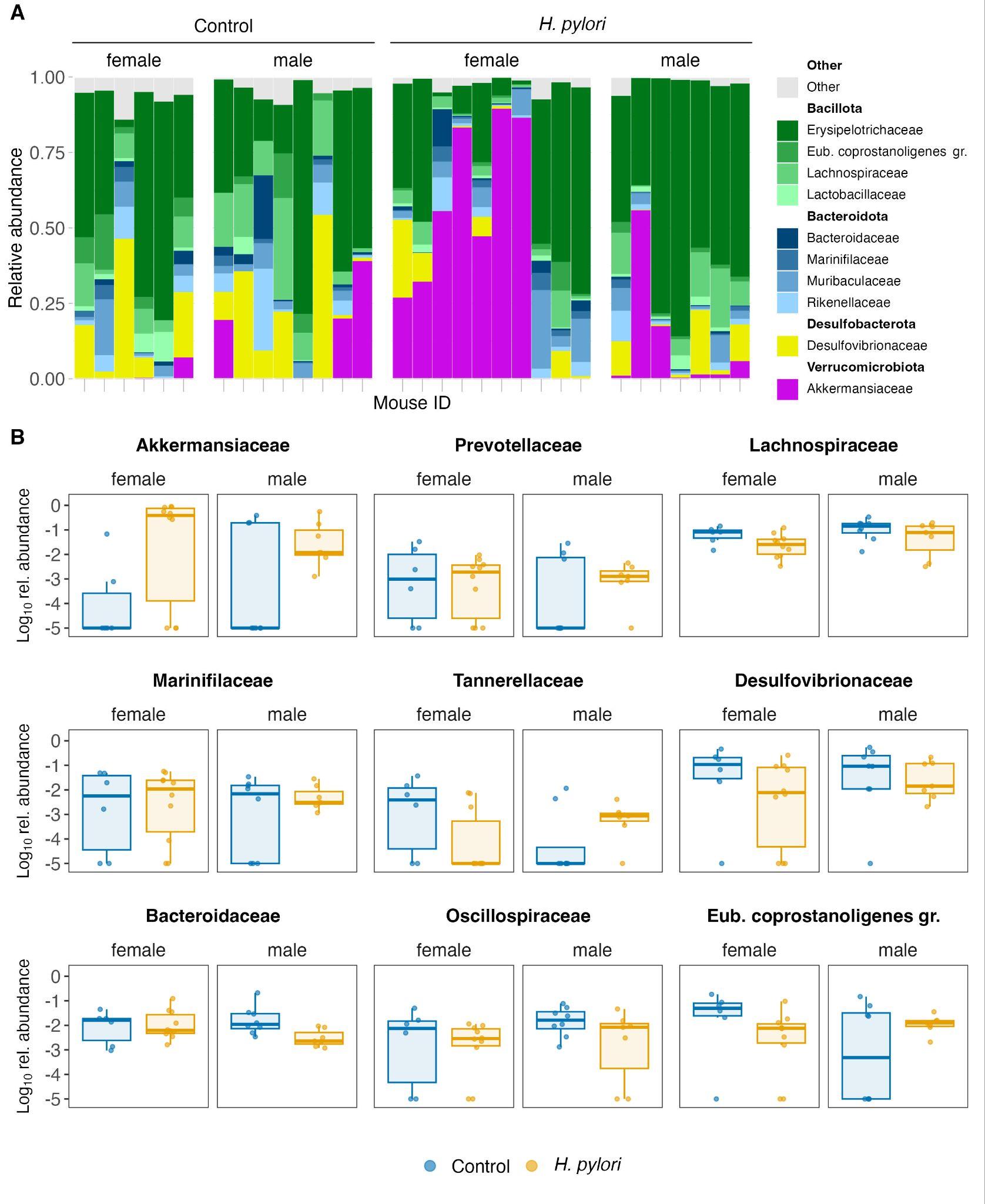

**Figure S15: Long-term experiment.** Colon microbiome composition in *H. pylori* infected or control C57BL/6JRj mice, measured by 16S rRNA gene sequencing. **A** Bars represent stacks of the 10 most relative abundant bacterial families in individual mice and are sorted in panels according to sex and treatment. The colors indicate the phylum, and the gradient of a given color represents different families within the phylum. **B** Nine colonic families, which showed differences in prevalence between *H. pylori*-infected (yellow) and control (blue) mice, stratified by sex. The relative abundances were log_10_-transformed prior to the analysis.

**SUPPLEMENTARY TABLES**

**Table S1: Short-term experiment.** Overview of the mice used in the short-term experiment, describing for each group defined by sex, treatment and diet the number of individuals, the litters they came from and the number of experimental cages.

| **Sex** | **Treatment** | **Diet** | **Individuals** | **Litters** | **Cages** |
| --- | --- | --- | --- | --- | --- |
| Males | Control | Control | 4 | 2, 1a, 5a | 1 |
| Females | Control | Control | 3 | 5a, 2 | 1 |
| Males | *H. pylori* | Control | 3 | 5b, 7, 4, 1b | 1 |
| Females | *H. pylori* | Control | 6 | 5b, 7, 4, 1b | 2 |
| Males | Control | HFD | 7 | 1a, 2, 5a | 2 |
| Females | Control | HFD | 5 | 5a, 2, 1a | 2 |
| Males | *H. pylori* | HFD | 5 | 4, 1b, 7 | 2 |
| Females | *H. pylori* | HFD | 9 | 7, 4, 5b, 1b | 3 |

**Table S2:** Samples contaminated with *Helicobacteraceae* reads, which were removed from subsequent analyses.

| mouse ID | experiment | type | treatment | weeks on diet | Helicobacteraceae (%) |
| --- | --- | --- | --- | --- | --- |
| EO4.11_FN | long-term | fecal | *H. pylori* | 0 | 0.2 |
| EO4.5_FL | long-term | cecal | *H. pylori* | 20 | 0.03 |
| EO4.10_ML | long-term | gastric | Control | 21 | 0.07 |
| EO5.12_MN | short-term | gastric | Control | 3 | 0.06 |

**SUPPLEMENTARY ONLINE METHODS**

**Isolation of stromal vascular cells (SVC) from perigonadal white adipose tissue (WAT)**

In brief, the tissue was minced and digested for 45 min at 37 °C in Hanks’ balanced salt solution with Ca^2+^ and Mg^2+^ supplemented with 0.5% bovine serum albumin (BSA, Sigma-Aldrich) and 10 mg/mL type II collagenase (Sigma-Aldrich), while rotating. EDTA was added [final concentration 10 mM] and the tissue further digested for 10 min. The cell solution was passed through a 100 μM nylon filter (Corning, Thermo) and then centrifuged at 4 °C for 10 min at 500 g. The cell pellet was lysed with 0.5 mL red blood cell lysis buffer (eBioscience) for 5 min before neutralization with 5 mL Flow Cytometry Staining (FACS) buffer (1% heat-inactivated FBS, 1 mM EDTA, 25 mM HEPES in 1X Phosphate-Buffered Saline (PBS)). The lysed cell solution was centrifuged as described above, and the cell pellet was dissolved in 100 μL FACS buffer.

**Cell counting and flow cytometry of SVCs from perigonadal WAT**

Briefly, cells were incubated 10 min with 0.5 μg/well Fc block (anti-CD16/CD32, 93 to reduce nonspecific antibody binding, then stained with 0.2 μL/well Fixable Viability Dye eFluor™ 506 in PBS for 30 min and a mix of the following antibodies at 0.2 μg/well: CD45.2 (104, PerCP-Cyanine5.5 conjugate); F4/80 (BM8, PE conjugate); CD11c (N418, PE-Cy7 conjugate); CD301 (ER-MP23, Alexa Fluor 647 conjugate); and 0.16 μg/well CD11b (M1/70, APC eFluor 780 conjugate) in FACS buffer for 20 min. All dyes were from eScience except ER-MP23 from BioRad. Cells were fixed with 2% paraformaldehyde for 10 min and acquired the following day on a CytoFlex flow cytometer (Beckman Coulter). Data were analysed with the FlowJo software (version 10.8.1). Live CD45⁺F4/80⁺CD11b⁺ macrophages were analyzed for CD11c and CD301 surface expression to identify macrophage subsets commonly referred to as M1-like (CD11c⁺) and M2-like (CD301⁺) in adipose tissue. Gates were established using fluorescence-minus-one (FMO) controls (Fig. S1).

***H. pylori* PMSS1 quantification**

For quantitative determination of *H. pylori* colonization, DNA was extracted from one quarter of the glandular stomach with the DNeasy Blood & Tissue Kit (Qiagen). For the DNA extraction, the following adaptations were made to the manufacturer’s protocol: (1) the tissue was lysed overnight at 56 °C and 300 rpm, and (2) DNA was eluted in 100 μL elution buffer. Infection status was first verified by conventional PCR (35 cycles) with primers targeting a region upstream of the the *OipA* gene: 5’-TCCCCGCGGGAAGTATCCAGGCGCTCCAT-3’ and 5’-GGACTAGTCGCCGATACTACCTTGTCCT-3’. Positive samples were analysed by absolute real-time PCR (qPCR) quantification with a standard curve, with a 20 μL reaction mix, consisting of 2 μL template, 0.8 μL of each forward and reverse primer (10 μM) and 10 μL of a 2x qPCRBIO SyGreen Mix (PCR Biosystems). Primers used in this study were: glmM F: 5’-GGATAAGCTTTTAGGGGTGTTAGGGG-3’ ^49^ and *GlmM* R: 5’-GCATTCACAAACTTATCCCCAATC-3’ ^50^. The samples and standards were run in duplicate with an initial denaturation at 95 °C for 3 min, followed by 40 cycles of 95 °C for 5 sec and 60°C for 30 sec. The standard curve was constructed using tenfold serial dilutions of a pGEM®-T Easy Vector with an inserted *glmM* gene. The concentration of the plasmid was converted to the number of *H. pylori* genome copies by the following equation: number of copies = (amount (ng) * 6.022×10^23^) / (plasmid length (bp) * 1×10^9^ * 650 Daltons). Based on the standard curve, the number of *H. pylori* copies in each sample was calculated with the following equation: number of copies = 10^Ct-Intercept^ / slope. Levels were standardized to the total DNA per sample as a proxy for host tissue amount, which was measured with a Qubit dsDNA high sensitivity assay kit on a Qubit 2.0 Fluorometer (Invitrogen).

**DNA extraction and 16S metabarcoding library preparation**

DNA from fecal pellets and content from ileum, cecum and colon was extracted in a randomized order with the ZymoBIOMICS Quick-DNA Fecal/Soil Microbe 96 Magbead Kit (Zymo Research) in a pre-PCR laboratory according to the manufacturer’s protocol. Four extraction controls were included on each 96-well plate to assess for potential contamination. DNA was eluted in 50 μL extraction buffer and stored at -20 °C until further processing. The V3-V4 (341F and 805R) region of the 16S rRNA gene was amplified with a broadly used primer set [40] containing Illumina adapter overhang sequences. The primers were:

5'-*TCGTCGGCAGCGTCAGATGTGTATAAGAGACAG*CCTACGGGNGGCWGCAG-3’ & 5' *GTCTCGTGGGCTCGGAGATGTGTATAAGAGACAG*GACTACHVGGGTATCTAATCC-3’ with the adapter sequence indicated in italics. A Microbial Community DNA Standard (Zymo Research) was included as a positive control. PCR amplifications were carried out on an Applied Biosystem 2720 Thermal Cycler with a reaction volume of 25 μL containing: 14.725 µl ddH2O, 2.5 µl (10X) Gold Buffer (GeneAmp®), 2.5 µl (25 mM) MgCl2, 0.625 µl (20 ng/µl) BSA, 0.5 dNTPs (10 mM), 0.15 µl (5 U/µl) DNA polymerase (AmpliTaq Gold®), 1 µl (10 mM) of each forward and reverse primer and 2 µl DNA extract. Two non-template controls were included per 96-well plate. The PCR program was set to 10 min at 95°C, 15 sec at 95°C, 20 sec at 55°C and 40 sec at 72°C, with a final elongation for 10 min at 72°C. To ensure that PCR inhibitors were not present and to find an appropriate cycle number, a subset of the samples was tested as undiluted, 1:10 and 1:100 diluted with qPCR before the PCR amplification. The PCR amplification was subsequently carried out with undiluted DNA and 25 cycles for the gut content extractions and 33 cycles for the gastric tissue extractions. The PCR products were shown on a 2% agarose gel by mixing 5 μL PCR product with 2 μL loading buffer. Amplicons were purified using magnetic beads (HighPrep™ PCR Clean-up System, MagBio Genomics Inc) following the manufacturer’s protocol, but with a bead:DNA ratio of 1:1. Library preparation was carried out with dual indexing using Nextera XT Index Kit Set A and Set D (Illumina) to reach 384 unique combinations. The index PCR was carried out with a total reaction volume of 28 μL containing 7 μL ddH20, 12 μL Accuprime Supermix II, 2 μL of each index primer and 5 μL PCR product. The PCR program was set to 2 min denaturation at 95 °C, 11 cycles of 15 sec at 95 °C, 20 sec of annealing at 55 °C and 40 sec elongation at 72 °C, with a final elongation for 10 min at 72 °C. The PCR products were visualized on an agarose gel as described above. Based on the band strength, amplicons were pooled (weak = 10 μL and strong = 5 μL) and purified with magnetic beads as described above. The final library was sequenced using an Illumina MiSeq platform.
